# TREM2-stimulated, PDGF-dependent macrophage-fibroblast crosstalk induces beneficial plaque remodeling in clonal hematopoiesis

**DOI:** 10.64898/2026.11.09.765508

**Authors:** Wenli Liu, Chenyi Xue, Malgorzata Olszewska, Cheng-Chieh Hsu, Mustafa Yalcinkaya, Tong Xiao, Kai Schlepckow, Christian Haass, Eirini P. Papapetrou, Kathryn M. Monroe, Nan Wang, Muredach P. Reilly, Alan R. Tall

## Abstract

**Background:** In *Jak2^V617F^* clonal hematopoiesis, inflammatory signaling suppresses macrophage TREM2 and promotes features of plaque instability, while therapeutic TREM2 agonism reverses plaque instability by increasing fibroblasts and fibrous cap formation. We tested whether macrophage-derived PDGFB and stromal cell PDGFRα signaling mediate this response.

**Methods:** Low allele burden (LAB) *Jak2^V617F^* atherosclerosis models were combined with inducible *Pdgfb* deletion in CX3CR1-lineage monocytes/macrophages or *Pdgfra* deletion in Col1a2-lineage fibroblast-enriched stromal cells. Mice were treated with the TREM2 agonist 4D9. Plaque composition, macrophage-fibroblast interactions, and aortic stromal cell populations were analyzed. Human WT and *JAK2^V617F^* iPSC-derived macrophages were used for validation.

**Results:** TREM2 activation with 4D9 increased macrophage PDGFB production and promoted fibrous cap formation in LAB *Jak2^V617F^* atherosclerosis. Deletion of *Pdgfb* in CX3CR1-lineage cells markedly attenuated the 4D9-mediated increase in fibrous cap area and lesional PDGFRα^+^ cells, establishing macrophage-derived PDGFB as a key signal for the stromal response. Consistently, Col1a2-lineage *Pdgfra* deletion abolished the 4D9-induced fibrous cap response without significantly altering lesion or necrotic core area. Single cell RNA-seq identified increased Fmod^+^ and CD26/Dpp4^+^ fibroblast populations associated with 4D9 treatment. Fmod^+^ fibroblasts represented an abundant matrix-associated population, whereas CD26^+^ fibroblasts showed greatest enrichment, prominent extracellular matrix, collagen organization, and matrix-remodeling programs that were enhanced by 4D9 and attenuated by *Pdgfra* deletion. The stromal response was also associated with reduced macrophage migration-related programs and predicted chemokine communication from CD26^+^ fibroblasts to macrophages which was verified by macrophage migration studies. Moreover, stromal *Pdgfra* deletion increased lesional CCR2^+^ and CCR5^+^ macrophages.

**Conclusions:** In a CH model of inflammatory atherosclerosis, TREM2 activation promotes a macrophage PDGFB-stromal PDGFRα pathway that expands Fmod^+^ and CD26^+^ fibroblast populations and increases fibrous cap matrix formation. CD26^+^ fibroblasts impair macrophage chemotaxis in a DPP4-dependent manner, suggesting reciprocal macrophage-fibroblast communication that coordinates matrix remodeling and inflammatory cell trafficking in the plaque.

**Graphical summary:** 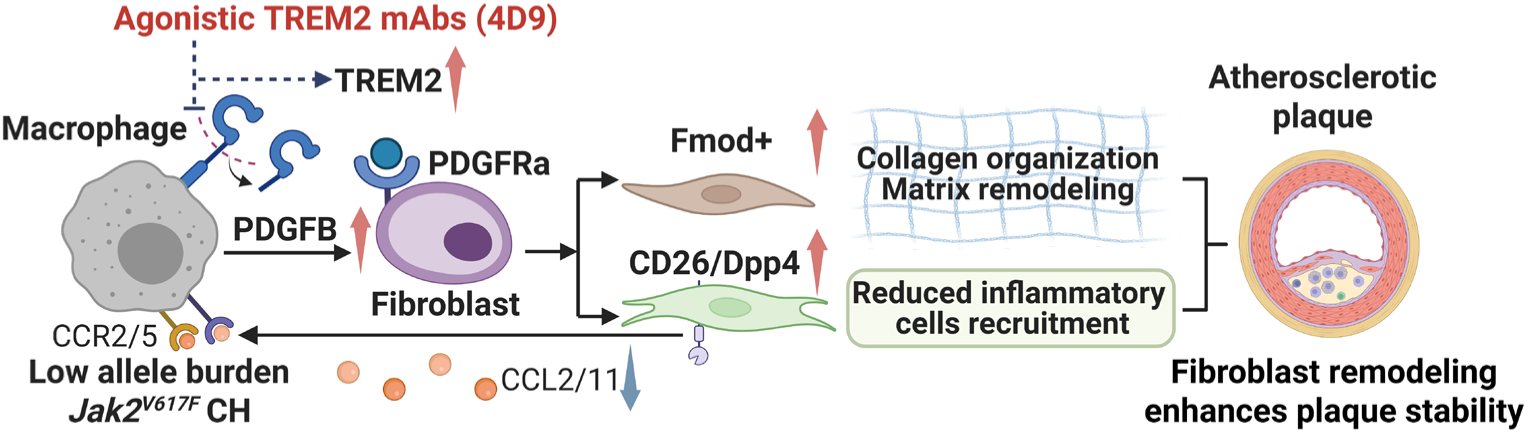

## Introduction

Cardiovascular disease remains the leading cause of death worldwide, with myocardial infarction and stroke accounting for most cardiovascular deaths^1,2^. Atherosclerotic plaque rupture and subsequent thrombosis are major causes of these acute cardiovascular events^3^. Although lesion size contributes to disease burden, plaque composition is an important determinant of clinical complications^4^. Stable plaques are characterized by an extracellular matrix-rich fibrous cap that separates the thrombogenic plaque core from the arterial lumen. While smooth muscle cells (SMC) have traditionally been considered the principal source of fibrous-cap matrix, lineage-tracing and single-cell studies have shown that SMCs can undergo phenotypic modulation into fibroblast-like “fibromyocytes” that populate the fibrous cap and promote features of plaque stability^5^. In addition to these SMC-derived cells, emerging evidence indicates that non-SMC derived vascular fibroblast populations can migrate into atherosclerotic lesions, contribute to extracellular matrix remodeling and fibrous cap formation, and promote features of plaque stability^6,7^. Lesional matrix-synthesizing fibroblast populations are also conserved between mice and humans and correlate with features of plaque stability features^8,9^. While Aortic fibroblasts remain understudied in atherosclerosis, and the signals that regulate their recruitment, expansion, and function within advanced plaques are poorly understood.

Within the atherosclerotic plaque, macrophages can regulate neighboring stromal cells through paracrine signaling. TREM2 is highly expressed in lipid-associated macrophage populations within atherosclerotic lesions and regulates macrophage lipid handling, survival and plaque remodeling^10,11^. TREM2 agonism has been proposed as a novel therapy to ameliorate Alzheimer’s Disease^12^ and to reduce cardiovascular risk^10,11^. In our previous study of low allele burden (LAB) *Jak2^V617F^* clonal hematopoiesis (CH), we found that inflammatory signaling reduced macrophage TREM2 and promoted features of plaque instability^13^. Conversely, treatment with the agonistic TREM2 antibody 4D9 restored TREM2 expression and increased fibrous cap formation in LAB mice^13^. These changes were particularly prominent in the CH setting reflecting increased macrophage inflammasome activation and IL-1β production^13^. Mechanistically, TREM2 activation increased PDGFB expression in macrophages through a SYK/AKT/β-catenin pathway and was associated with accumulation of PDGFRα-positive fibroblast-like cells within the fibrous cap^13^. These findings suggested that in the inflammatory setting of LAB CH where macrophage TREM2 expression is reduced, TREM2 activation may promote fibrous cap formation through a macrophage PDGFB-fibroblast PDGFRα signaling axis. However, whether macrophage-derived PDGFB and fibroblast PDGFRα signaling are causally required for this response remains unknown.

PDGF signaling has complex and context-dependent roles in atherosclerosis. PDGF has traditionally been associated with vascular mesenchymal cell proliferation and migration, and previous studies have implicated PDGF signaling in both lesion growth and extracellular matrix accumulation^14,15^. However, hematopoietic *Pdgfb* deficiency delays fibrous cap formation, while alterations in PDGF signaling can influence plaque collagen content and the abundance of mesenchymal cells without necessarily producing parallel changes in lesion size^16,17^. The effects of PDGF signaling are receptor and cell type dependent. Functional blockade of PDGFRβ markedly reduced vascular SMC accumulation within fibrous lesions in *Apoe^−/−^* mice^18^. Consistently, constitutive activation of PDGFRβ in SMCs induced chemokine expression and vascular inflammation and markedly accelerated atherosclerosis in hypercholesterolemic mice^19^. More recently, genetic studies of the coronary artery disease-associated PDGFD locus further linked PDGF signaling to changes in vascular SMC expansion and phenotypic state as well as inflammatory responses in adventitial mesenchymal cells, without corresponding changes in overall plaque burden^20^. In contrast to the well-established role of PDGFRβ in VSMCs, the function of PDGFRα signaling in atherosclerosis remains poorly defined. PDGFRα is enriched in arterial fibroblasts in both mice and humans, including fibroblast populations associated with collagen organization and vascular remodeling^9^. Thus, whether PDGFRα identifies a functionally distinct fibroblast response to macrophage-derived PDGFB in advanced atherosclerosis, and whether this signaling contributes to fibrous cap formation remains unknown.

Here, we investigated the cellular and molecular mechanisms by which TREM2-induced macrophage PDGFB signaling regulates fibroblast responses and fibrous cap formation. Using complementary conditional deletion of *Pdgfb* in CX3CR1-lineage monocytes/macrophages and *Pdgfra* in Col1a2-lineage stromal cells, together with macrophage-fibroblast conditioned-media studies, we establish that macrophage-derived PDGFB and stromal PDGFRα are required for the fibrous cap response to TREM2 activation in LAB mice. Our single-cell analysis of aortic stromal cells similarly identified heterogeneous fibroblast populations, including a CD26^+^ PDGFRα-expressing population enriched for extracellular matrix and collagen-associated gene expression programs and an Fmod^+^ fibroblast population with a distinct matrix-remodeling program. TREM2 activation preferentially expanded the CD26+ fibroblast population and enhanced its matrix-associated phenotype through PDGFRα signaling while also suppressing fibroblast to macrophage back-talk. Together, these findings define a TREM2-PDGFB-PDGFRα pathway that coordinates macrophage and fibroblast responses during fibrous cap formation and identify CD26^+^ fibroblasts as both a matrix-associated and immunoregulatory stromal population in atherosclerosis.

## Results

### Monocyte/macrophage-derived PDGFB is required for 4D9-induced fibrous cap formation

We previously showed that activation of TREM2 with the agonistic antibody 4D9 increased fibrous cap area in LAB *Jak2^V617F^* atherosclerosis in association with increased macrophage *Pdgfb* expression and predicted macrophage PDGFB-fibroblast PDGFR signaling^13^. Because fibrous cap formation is a key feature of plaque stabilization, and prior studies have implicated PDGF signaling in stromal cells and fibrous cap responses in atherosclerosis, we tested whether macrophage-derived PDGFB is required for the plaque-stabilizing effect of TREM2 activation.

Given that TREM2 agonism appeared to act primarily on the major population of non-mutant macrophages^13^, we generated LAB *Jak2^V617F^* bone marrow (BM) chimeric mice in which the non-mutant myeloid compartment was either PDGFB sufficient or deficient (Fig. 1A). Female *Ldlr^−/−^* recipient mice were reconstituted with 1.5% *Jak2^V617F^* GFP^+^ BM together with 98.5% *Pdgfb^fl/fl^* or 98.5% littermate control *Pdgfb^fl/fl^ Cx3cr1CreER* BM, referred to as LAB and LAB/PDKO mice, respectively (Fig. 1A). Tamoxifen treatment was initiated 4 weeks after BM transplantation and maintained throughout the study to sustain *Pdgfb* deletion in continuously replenished CX3CR1-expressing macrophages. Mice were then fed Western diet and treated with IgG or 4D9 for 14 weeks (Fig. 1A). Blood counts, body weight, and spleen/body weight ratio were comparable between groups, indicating that monocyte/macrophage *Pdgfb* deletion did not cause major systemic or hematologic changes (Supplemental Fig. 1A-B).

**Fig 1.**
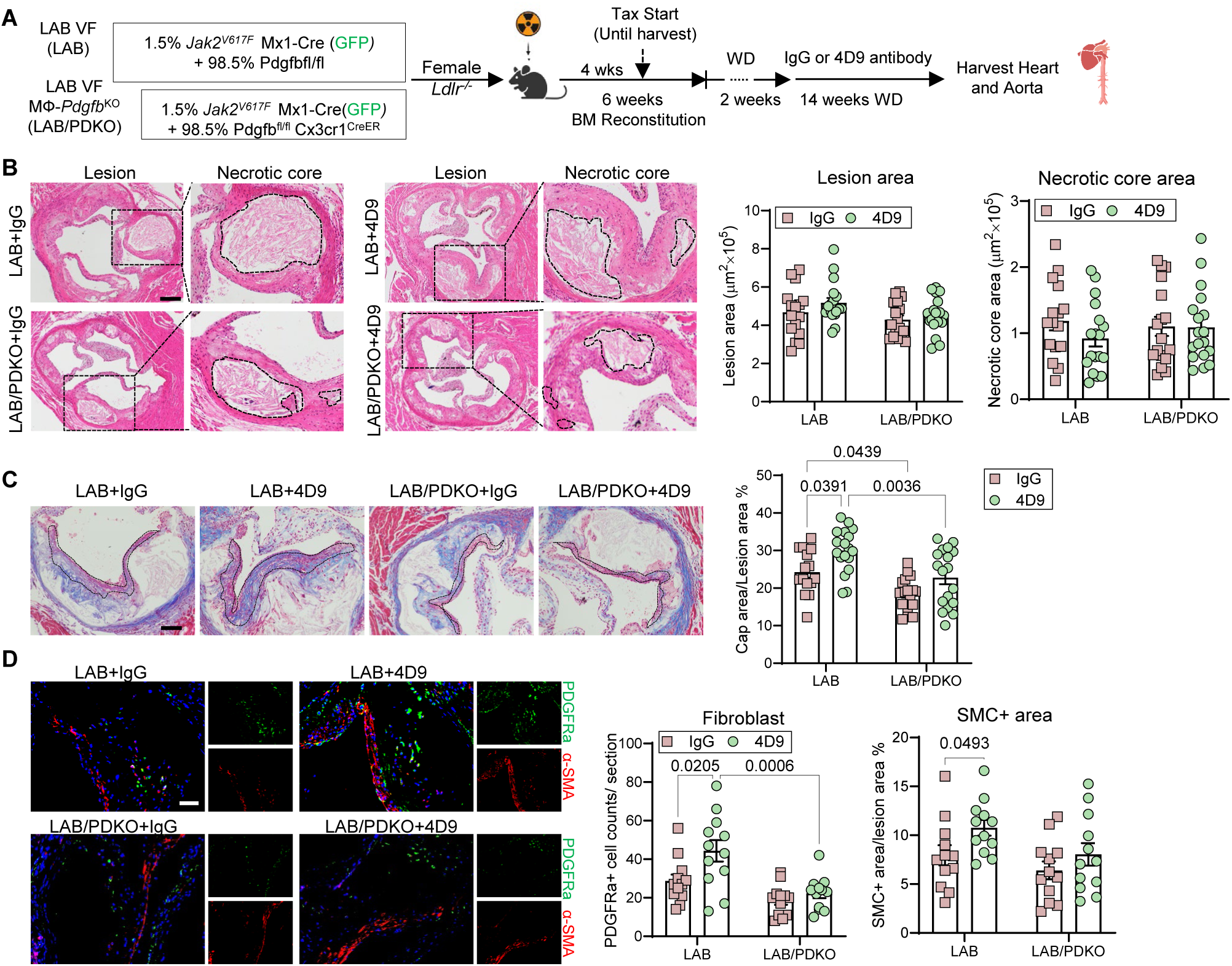
PDGFB deficiency reverses the beneficial effect of 4D9 on cap area by reducing fibroblast and smooth muscle cell accumulation. **A.** Experimental design. The mice received oral tamoxifen treatment 2-3 times per week, starting 4 weeks after BMT and continuing until the end of the study. **B.** Representative H&E staining and quantification of lesion area and necrotic core area in the aortic root sections. Necrotic core regions indicated by broken lines. Scale bar, 200μm. **C.** Aortic root sections were stained with Masson’s trichrome for fibrous cap and quantified as the ratio of total lesion area. Scale bar, 75μm. **D.** Representative immunofluorescence staining of PDGFRa (fibroblast) and α-SMA (smooth muscle cells) and quantification of lesional fibroblast numbers and α-SMA+ area. Scale bar, 50μm.

Macrophage *Pdgfb* deletion did not substantially alter total lesion area or necrotic core area after IgG or 4D9 treatment (Fig. 1B). In contrast, Masson’s trichrome staining showed that 4D9 significantly increased fibrous cap area in LAB mice, whereas this response was lost in LAB/PDKO mice (Fig. 1C), demonstrating that macrophage-derived PDGFB is required for the fibrous cap response to TREM2 activation (Fig. 1C). PDKO also reduced fibrous cap area in IgG control mice consistent with a role of PDGFB in fibrous cap formation under basal conditions. We next examined the cellular changes associated with this impaired cap response. 4D9 increased lesional PDGFRα^+^ fibroblast accumulation in LAB mice, but this increase was abolished by macrophage *Pdgfb* deletion. Lesional α-SMA^+^ area showed similar though less pronounced changes, increasing after 4D9 treatment in LAB mice but not significantly in LAB/PDKO mice (Fig. 1D). Together, these data show that macrophage-derived PDGFB is necessary for 4D9-induced fibroblast accumulation and increased fibrous cap area in LAB *Jak2^V617F^* atherosclerosis.

### Macrophage-derived PDGFB supports fibroblast proliferation and migration following 4D9 treatment

We next asked whether macrophage-derived PDGFB acts directly through fibroblast PDGF receptors to regulate aortic fibroblast function. BM-derived macrophage (BMDMs) from LAB and LAB/PDKO mice from the above cohort were treated with IgG or 4D9 for 48 hours, and macrophage-conditioned media were collected (Fig. 2A). To examine the response of primary aortic fibroblasts, CD45^-^CD90.2^+^αSMA^-^ fibroblasts were isolated by flow cytometry from WT mice aortas, separating them from αSMA^+^ SMCs and CD45^-^CD90.2^+^αSMA^+^ myofibroblasts, and subsequently exposed to the indicated macrophage-conditioned media (Fig. 2A). Fibroblasts were additionally treated with or without the PDGFR inhibitor CP673451 (CP), followed by assessment of proliferation and migration (Fig. 2A). Consistent with the in vivo findings, 4D9 increased PDGFB abundance in conditioned media from LAB macrophages, whereas PDGFB levels were markedly reduced in conditioned media from LAB/PDKO macrophages regardless of antibody treatment, confirming the effect of 4D9 on PDGFB and the knockout efficiency (Fig. 2B). We assessed fibroblast proliferation by EdU incorporation. Conditioned media from 4D9-treated LAB macrophages increased fibroblast proliferation compared with media from IgG-treated LAB macrophages. This proliferative response was attenuated when fibroblasts were exposed to conditioned media from LAB/PDKO macrophages or treated with the PDGFR inhibitor CP (Fig. 2C). These findings indicate that macrophage-derived PDGFB and fibroblast PDGFR signaling contribute directly to fibroblast proliferation in the setting of TREM2 activation.

**Fig 2.**
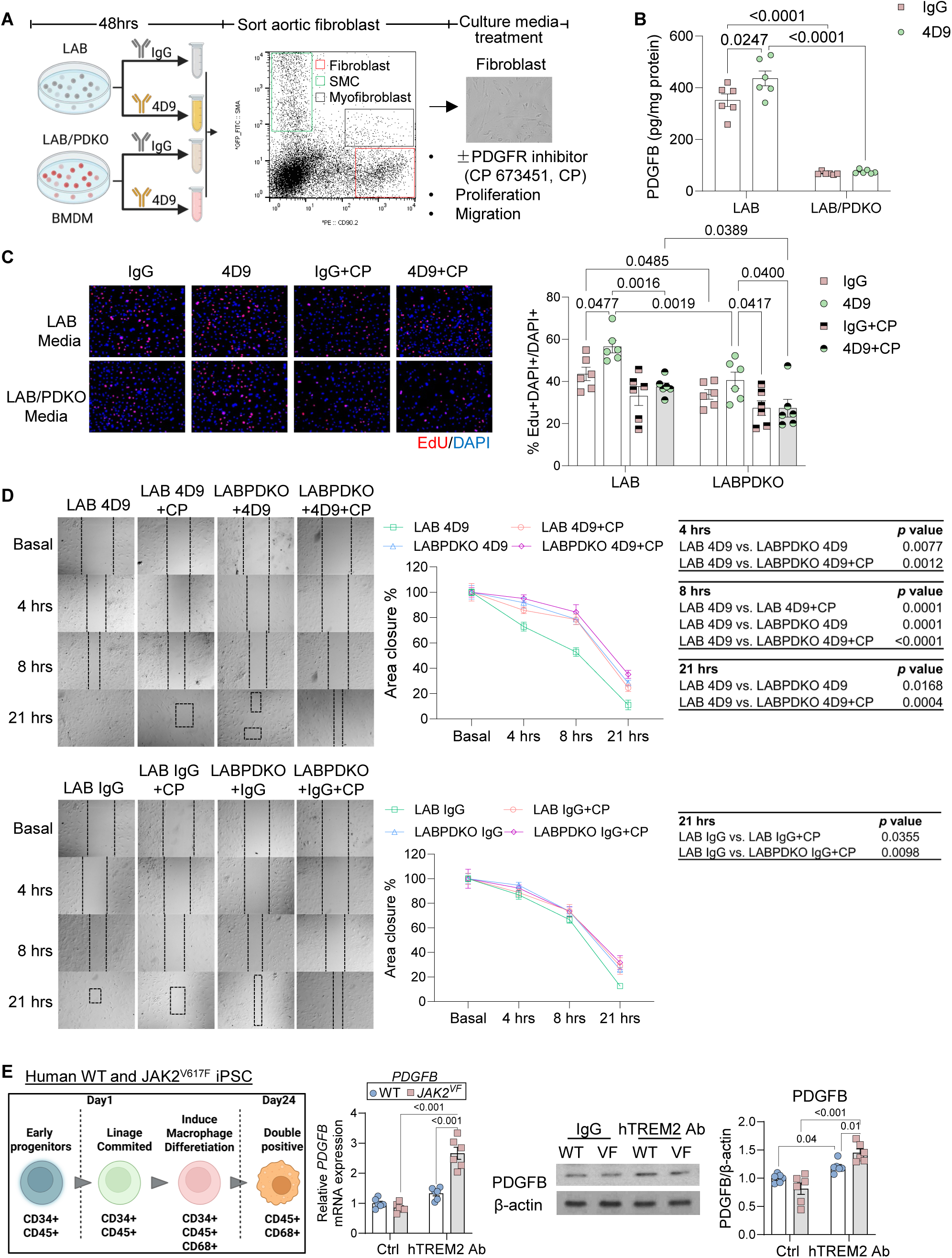
Macrophage-derived PDGFB promotes aortic fibroblast proliferation and migration through PDGFR signaling. **A.** Experimental workflow for macrophage-conditioned media studies. BMDMs from LAB and LAB/PDKO mice were treated with IgG or 4D9 (20μg/ml) for 48 hours, and conditioned media were collected and applied to sorted aortic fibroblast in the presence or absence of the PDGFR inhibitor CP-673451 (CP) for assessment of proliferation and migration. **B.** PDGFB abundance in conditioned media from LAB and LAB/PDKO BMDMs treated with IgG or 4D9. **C.** Representative EdU staining and quantification of EdU+ fibroblast following treatment with conditioned media from the indicated groups, with or without CP-673451 (100nM). **D.** Representative scratch-wound images and quantification of fibroblast migration following treatment with conditioned media from the indicated groups, with or without CP-673451 (100nM). Fibroblasts were pretreated with mitomycin C (5 μg/mL) for 2 hours before the assays. Wound closure was assessed at baseline and 4, 8, and 21 hours. Results for conditioned media from 4D9-treated macrophages are shown above and those from IgG-treated macrophages below. **E.** Schematic of human iPSC-derived macrophage differentiation. Isogenic WT and *JAK2^V617F^* iPSC-derived macrophages were treated with IgG or an agonistic human TREM2 antibody (20μg/ml) for 48 hours. PDGFB mRNA expression was assessed by qPCR, and PDGFB protein abundance was assessed by immunoblotting with β-actin as a loading control.

We then examined fibroblast migration using a scratch assay. To minimize the contribution of proliferation to wound closure, fibroblasts were pretreated with proliferation inhibitor mitomycin C. Conditioned media from 4D9-treated LAB macrophages produced more rapid wound closure than media from 4D9-treated LAB/PDKO macrophages, with differences detectable as early as 4 hours and maintained at 8 and 21 hours (Fig. 2D, upper panel). Pharmacological inhibition of PDGFR signaling also reduced the migratory response to LAB 4D9-conditioned media, with a clear effect at 8 hours, while the combination of macrophage *Pdgfb* deficiency and PDGFR inhibition produced the greatest impairment in wound closure (Fig. 2D). Under IgG-treated conditions, PDGFR inhibition only modestly reduced wound closure at 21 hours (Fig. 2D, lower panel). These findings indicate that macrophage-derived PDGFB and fibroblast PDGFR signaling contribute to the migratory response induced by TREM2 activation.

To determine whether TREM2-dependent induction of macrophage PDGFB was conserved in human cells, we examined isogenic WT and *JAK2^V617F^* human iPSC-derived macrophages. Treatment with an agonistic human TREM2 antibody increased *PDGFB* mRNA expression in both WT and *JAK2^V617F^* macrophages, but the response was greater and only significant in *JAK2^V617F^* cells (Fig. 2E). PDGFB protein abundance was significantly increased in both WT and *JAK2V^617F^* macrophages following TREM2 activation, with a greater response in antibody-treated *JAK2^V617F^* macrophages. These findings extend the mouse LAB BMDM observations to human macrophages and support a conserved effect of TREM2 activation on macrophage PDGFB expression.

Together, our data show that TREM2 activation increases macrophage-derived PDGFB and that this response promotes aortic fibroblast proliferation and migration through PDGFR-dependent signaling. These functional findings provide a direct mechanistic link between macrophage PDGFB induction and the increased fibroblast accumulation and fibrous cap remodeling observed following 4D9 treatment in vivo.

### Fibroblast PDGFRα is required for the fibrous cap response to 4D9

PDGFB can signal through both PDGFRα and PDGFRβ, which are expressed across vascular mesenchymal populations^21^. However, PDGFRα is enriched in arterial fibroblasts and shows limited overlap with differentiated SMC markers, whereas PDGFRβ has been more closely associated with SMC and myofibroblasts within atherosclerotic aorta^22^. We therefore next assessed whether the fibroblast-enriched PDGFRα was required for the 4D9-induced fibrous cap response.

To address this question in vivo, we crossed *Pdgfra*^fl/fl^ mice with Col1a2-CreERT mice on the *Ldlr*^-/-^ background. *Pdgfra*^fl/fl^*Ldlr*^-/-^ and *Pdgfra*^fl/fl^ Col1a2-CreERT *Ldlr*^-/-^ recipients were reconstituted with the same LAB *Jak2^V617F^* BM mixture, generating LAB and LAB/RaKO mice, respectively (Fig. 3A). Tamoxifen was administered after BM transplantation to induce *Pdgfra* deletion in Col1a2-lineage recipient stromal cells. Following BM reconstitution, mice were fed Western diet for 16 weeks, with IgG or 4D9 treatment during the final 14 weeks. Efficient reduction of PDGFRα in aortic CD90.2^+^ fibroblast-enriched stromal cells was confirmed by flow cytometry (Supplemental Fig. 2A). Blood counts, body weight, and spleen weight ratio were not altered by stromal *Pdgfra* deletion (Supplemental Fig. 2B-C).

**Fig 3.**
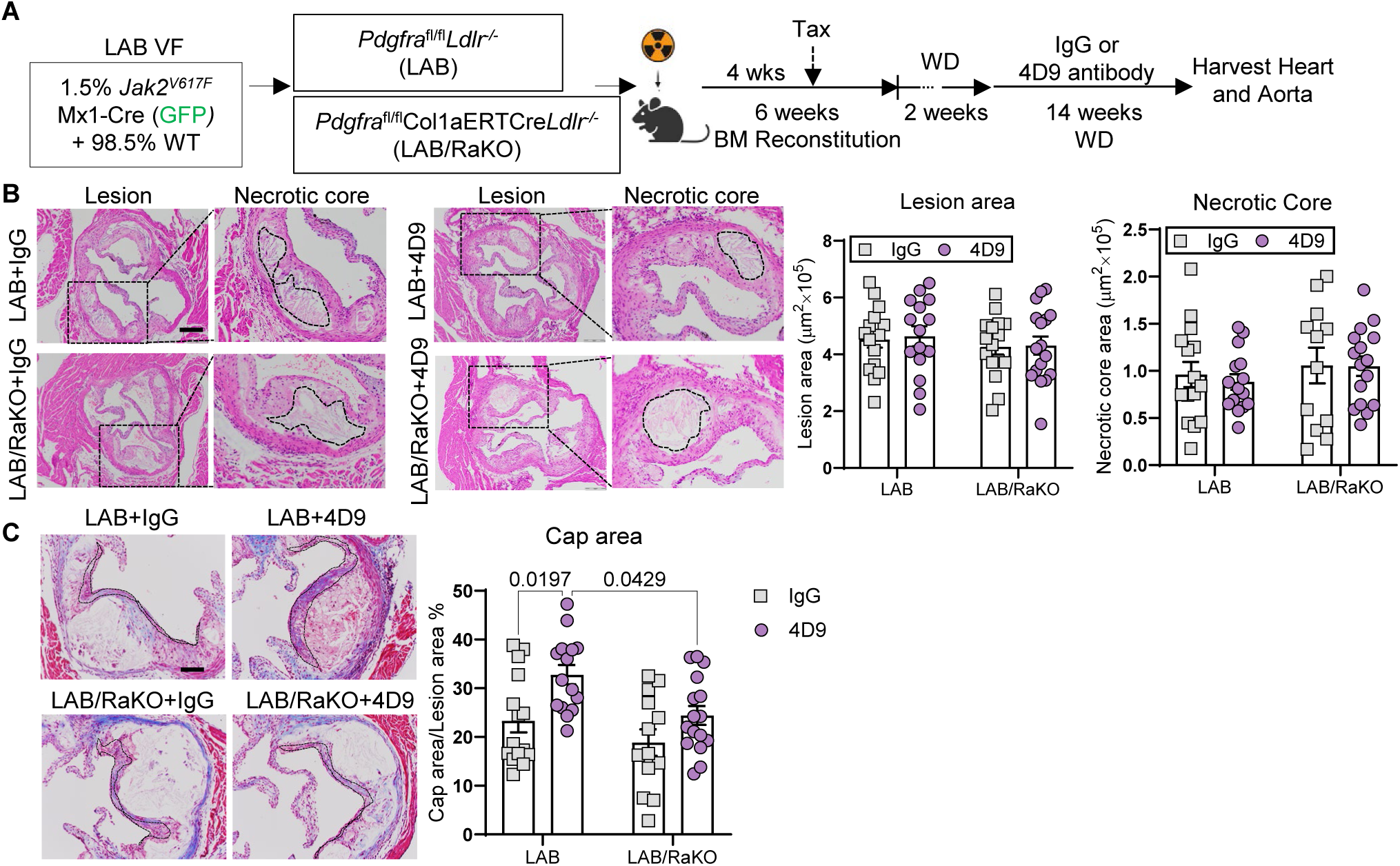
PDGFRa deficiency reverses the beneficial effect of 4D9 on fibrous cap area. **A.** Experimental design. **B.** Representative H&E staining and quantification of lesion area and necrotic core area in the aortic root sections. Necrotic core regions indicated by broken lines. Scale bar, 200μm. **C.** Aortic root sections were stained with Masson’s trichrome for fibrous cap and quantified as the ratio of total lesion area. Scale bar, 75μm. Scale bar, 50μm.

Aortic root analysis showed that stromal *Pdgfra* deletion did not significantly alter overall lesion area or necrotic core area under either IgG or 4D9-treated conditions (Fig. 3B). In contrast, Masson’s trichrome staining revealed a selective effect on fibrous cap formation (Fig. 3C). 4D9 increased cap area relative to lesion area in LAB mice as predicted, whereas this increase was significantly attenuated in LAB/RaKO mice (Fig. 3C). Moreover, cap area in 4D9-treated LAB/RaKO mice was significantly lower than in 4D9-treated LAB mice (Fig. 3C). Thus, loss of stromal PDGFRα selectively impaired the fibrous cap response to TREM2 activation without substantially affecting overall plaque burden or necrotic core size in LAB *Jak2^V617F^* CH mice. Together, these findings identify a stromal PDGFRα signaling axis that is required for the fibrous cap remodeling induced by 4D9.

### 4D9 reshapes fibroblast states and matrix-associated programs in a PDGFRα-dependent manner

To define the fibroblast populations underlying the PDGFRα-dependent fibrous cap response, we performed parallel single-cell RNA sequencing (scRNA-seq) of enriched aortic stromal and immune cell populations from LAB and LAB/RaKO mice treated with IgG or 4D9. For the stromal dataset, CD45^-^ αSMC^-^CD90.2^+^ cells were enriched to capture fibroblasts, while small fractions of αSMA^+^ vascular cells and CD45^+^ immune cells were intentionally retained to provide internal reference populations within the aortic microenvironment. Approximately 5% of the recovered cells were SMC and 30% were CD45^+^ immune cells. Unsupervised clustering identified seven stromal populations, including two fibromodulin^+^ (Fmod^+^) fibroblast populations, a CD26^+^ fibroblast population, a fibroblast 3 population, two SMC populations, and pericytes (Fig. 4A). Expression of representative fibroblast, SMC, and pericyte markers supported these annotations (Supplemental Fig. 3).

**Fig. 4.**
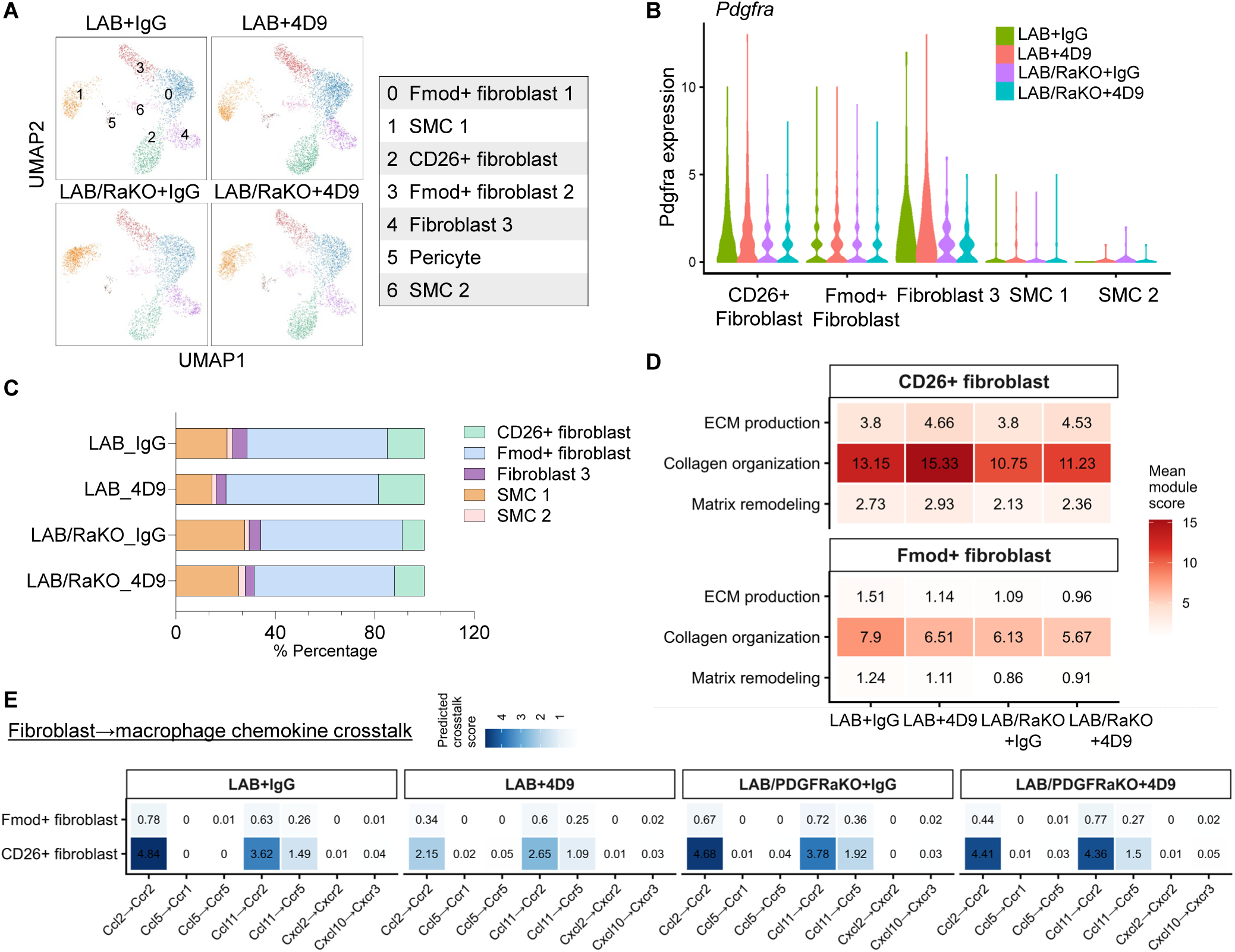
TREM2 activation and fibroblast PDGFRα deficiency modify fibroblast heterogeneity, matrix programs, and fibroblast-macrophage chemokine crosstalk. **A.** UMAP visualization of CD90.2⁺ aortic stromal cells from LAB and LAB/PDGFRaKO (LAB/RaKO) mice treated with IgG or 4D9 antibody. Clusters were annotated as Fmod⁺ fibroblast 1, SMC 1, CD26⁺ fibroblast, Fmod⁺ fibroblast 2, fibroblast 3, pericyte, and SMC 2. **B.** Violin plots showing log-normalized single-cell Pdgfra expression across fibroblast and SMC subpopulations. **C.** Relative abundance of major fibroblast and SMC populations from LAB and LAB/RaKO aortas treated with IgG or 4D9, based on single-cell RNA-sequencing data. **D.** Module-wise scaled scores for matrix remodeling, collagen organization, and ECM production programs in CD26⁺ fibroblasts and Fmod⁺ fibroblasts. **E.** Predicted fibroblast to macrophage chemokine signaling inferred from CellChat analysis, showing selected ligand-receptor interactions from Fmod⁺ and CD26⁺ fibroblasts to macrophages under IgG or 4D9 treatment in LAB and LAB/RaKO aortic cells. Color intensity indicates predicted communication strength. Data are shown as group-level single cell analyses.

Comparing PDGF receptor expression across these stromal populations. *Pdgfra* was preferentially expressed in the fibroblast populations, including CD26^+^ and Fmod^+^ fibroblast, whereas expression was substantially lower in the SMC populations (Fig. 4B). In contrast, *Pdgfrb* showed broader expression across both fibroblast and SMC populations (Supplemental Fig. 4A). *Pdgfra* expression was reduced in LAB/RaKO stromal cells, consistent with targeted deletion (Fig. 4B). These findings further supported PDGFRα as the fibroblast-enriched receptor through which macrophage-derived PDGFB could regulate the stromal response. scRNA-seq showed greater recovery of both Fmod^+^ and CD26^+^ fibroblasts following 4D9 treatment in LAB mice, whereas these increases were markedly attenuated in LAB/RaKO mice (Supplemental Fig. 4B). Analysis of relative cell composition showed that Fmod^+^ fibroblasts remained a similar fraction of the stromal compartment, whereas CD26^+^ fibroblasts showed increased fractional enrichment following 4D9 treatment (Fig. 4C). In contrast, Fibroblast 3 represented only a small fraction of the stromal compartment and showed no consistent change across treatment or genotype (Fig. 4C, Supplemental Fig 4B).

We next used an unbiased Gene Ontology (GO) enrichment analysis to define the fibroblast programs altered by 4D9 treatment. In CD26^+^ fibroblasts, genes increased by 4D9 were enriched among the top GO biological processes for extracellular matrix organization, extracellular structure organization, external encapsulating structure organization, and vascular morphogenesis/development (Supplemental Fig. 4C). In contrast, no significantly enriched GO biological processes were identified among genes increased by 4D9 in Fmod^+^ fibroblasts. We then asked whether these responses depended on stromal PDGFRα signaling. In 4D9-treated LAB/RaKO mice, extracellular matrix and extracellular structure organization were among the most prominently reduced GO terms in CD26^+^ fibroblasts compared with 4D9-treated LAB mice, and related matrix programs were also reduced in Fmod^+^ fibroblasts (Supplemental Fig. 4D). Consistent with the GO analysis, module-wise scaled scores showed that CD26^+^ fibroblasts had higher collagen organization, extracellular matrix (ECM) production, and matrix remodeling programs than Fmod^+^ fibroblasts (Fig. 4D). These programs were increased by 4D9 and attenuated following *Pdgfra* deletion. Individual matrix and collagen-associated genes showed a similar pattern, with stronger expression in CD26^+^ fibroblasts and a broad reduction after stromal *Pdgfra* deletion (Supplemental Fig. 4E). Together, these findings suggest that while both the more abundant Fmod^+^ fibroblasts and the less abundant CD26^+^ fibroblasts are supporting matrix programs in response to 4D9, the most dramatic changes occur in the CD26^+^ population and are dependent on *Pdgfra*.

### PDGFRα-dependent fibroblast remodeling is associated with macrophage migration programs

To determine how TREM2 agonist 4D9 altered plaque macrophage states and whether these changes were influenced by stromal PDGFRα signaling, we analyzed the CD45^+^ myeloid cells retained in the same aortic single-cell dataset. Unsupervised clustering identified two macrophage populations (Macrophage 1 and 2), a dendritic cell population, and neutrophils (Supplemental Fig. 5A). Two macrophage populations were identified by CD68 expression and showed expression of *Trem2* and lipid-associated macrophage genes including *Apoe*, *Lpl*, *Ctsd*, *Cd9* and *Lgals3*, together with *C1qa*, *C1qb*, *Adgre1* and *Lyz2* (Supplemental Fig. 5B-C). Macrophage 1 represented a prominent *Trem2*-expressing population and contained sufficient cells for differential-expression analysis. GO enrichment analysis showed that positive regulation of cell migration was among the biological processes downregulated by 4D9 in Macrophage 1 (Supplemental Fig. 5D). In contrast, cell migration was enriched among genes increased in LAB/RaKO compared with LAB Macrophage 1 under 4D9 treatment (Supplemental Fig. 5E), indicating that disruption of stromal PDGFRα signaling was associated with reversal of the diminished migration-related transcriptional response to 4D9.

For analysis of fibroblast-macrophage communication, Macrophage 1 and Macrophage 2 were combined to capture the broader macrophage compartment. CellChat analysis identified CD26^+^ fibroblasts as the predominant fibroblast source of predicted macrophage-directed chemokine signaling, with substantially higher *Ccl2/Ccl11-Ccr2* and *Ccl11-Ccr5* interaction scores than Fmod^+^ fibroblasts across all groups (Fig. 4E). In LAB mice, 4D9 reduced these predicted interactions, whereas stromal *Pdgfra* deletion increased *Ccl2/Ccl11-Ccr2* and *Ccl11-Ccr5* signaling under 4D9 treatment (Fig. 4E). These findings suggest that PDGFRα-dependent changes in the fibroblast compartment are accompanied by decreased fibroblast to macrophage chemokine communication. Consistent with the CellChat analysis, CD26^+^ fibroblasts expressed substantially higher levels of *Ccl2* and *Ccl11* than Fmod^+^ fibroblasts (Supplemental Fig. 5F). In LAB mice, 4D9 reduced *Ccl2* expression in CD26^+^ fibroblasts, whereas *Ccl2* remained high and *Ccl11* was further increased after stromal *Pdgfra* deletion (Supplemental Fig. 5F). Among macrophages, *Ccr2* was the most highly expressed chemokine receptor, with lower but detectable *Ccr5* expression (Supplemental Fig. 5G). These findings identified fibroblast to macrophage chemokine signaling as a potential link between the PDGFRα-dependent stromal response and macrophage migration.

### Macrophage PDGFB-stromal PDGFRα signaling promotes fibroblast-associated matrix remodeling

The single-cell analysis identified distinct Fmod^+^ and CD26^+^ fibroblast populations associated with matrix remodeling. We therefore examined these populations directly in atherosclerotic lesions. Picrosirius red staining showed a trend toward increased plaque collagen content following 4D9 treatment in LAB mice, whereas collagen content was reduced in LAB/RaKO mice under both IgG and 4D9 treatment conditions (Fig. 5A and Supplemental Fig. 6A).

**Fig 5.**
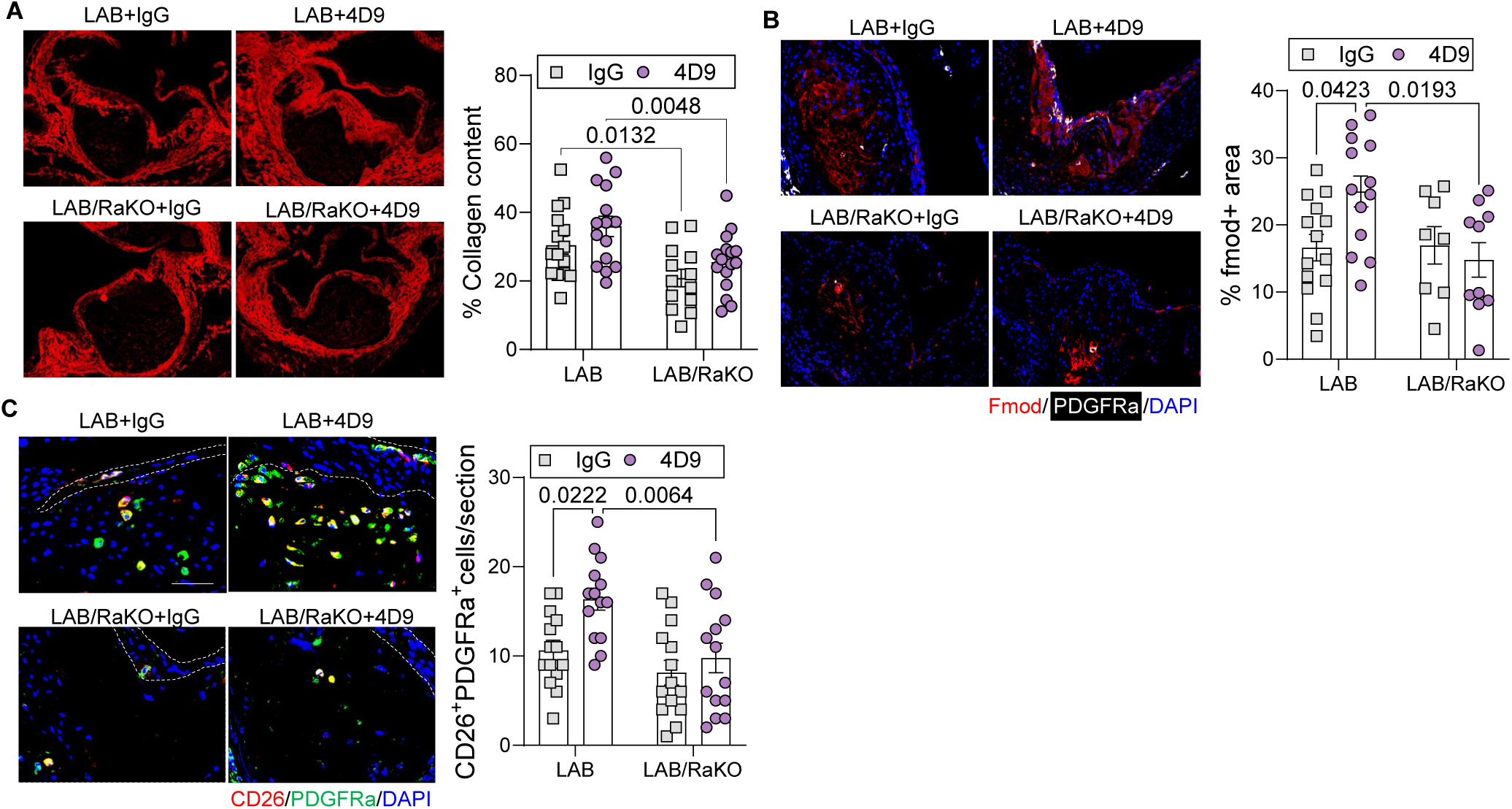
PDGFB-PDGFRα signaling regulates fibroblast populations and collagen accumulation in atherosclerotic plaques. **A.** Representative fluorescence images of Picrosirius red-stained aortic root lesions from LAB and LAB/RaKO mice treated with IgG or 4D9, with quantification of collagen content within the lesion as the percentage of the total lesion area. **B.** Representative immunofluorescence staining of aortic root lesions for Fmod (red), PDGFRα (white), and DAPI (blue), with quantification of Fmod^+^ area within the lesion. **C.** Representative immunofluorescence staining for CD26 (red), PDGFRα (green), and DAPI (blue). Dashed lines indicate the fibrous cap region, and arrowheads indicate CD26^+^PDGFRα^+^ cells. Right, quantification of CD26^+^PDGFRα^+^ fibroblast per section.

*Fmod* was expressed by an abundant fibroblast population in the single-cell dataset. Fibromodulin is a secreted extracellular matrix proteoglycan that regulates collagen fibril organization and is present within human and mouse atherosclerotic plaque matrix^23,24^. Consistent with its extracellular localization, Fmod staining showed a diffuse matrix-associated pattern within the lesion rather than discrete cellular staining (Fig. 5B). Fmod^+^ area increased following 4D9 treatment in LAB plaques and was reduced in LAB/RaKO mice (Fig. 5B), supporting PDGFRα-dependent remodeling of the Fmod-associated matrix compartment. Lesional CD26^+^ fibroblasts co-expressed PDGFRα and were readily identified as individual CD26^+^ fibroblast. 4D9 increased CD26^+^ fibroblasts in LAB plaques, whereas their accumulation was reduced following stromal *Pdgfra* deletion (Fig. 5C). The same response was independently examined in the LAB/PDKO cohort, where macrophage *Pdgfb* deletion reduced the 4D9-induced accumulation of CD26^+^ fibroblasts (Supplemental Fig. 6B). These findings place the CD26^+^ fibroblast response downstream of macrophage PDGFB-stromal PDGFRα signaling.

### CD26^+^ fibroblasts restrain macrophage recruitment through reciprocal chemokine signaling

The predicted fibroblast to macrophage communication identified by single-cell analysis led us to examine whether CD26^+^ fibroblasts could functionally regulate macrophage recruitment. Flow cytometric analysis of aortic CD45^-^α-SMC^-^CD90.2^+^ fibroblasts confirmed an increased proportion of CD26^+^ fibroblasts following 4D9 treatment in LAB mice (Fig. 6A and B). Because CD26 is encoded by *Dpp4* and functions as a cell-surface dipeptidyl peptidase, we next measured its enzymatic activity. Sorted CD26^+^ aortic fibroblasts displayed markedly greater DPP4 activity than either CD26^−^ fibroblasts or SMCs (Fig. 6C), establishing CD26 enzymatic activity as a functional feature of this fibroblast population.

**Fig 6.**
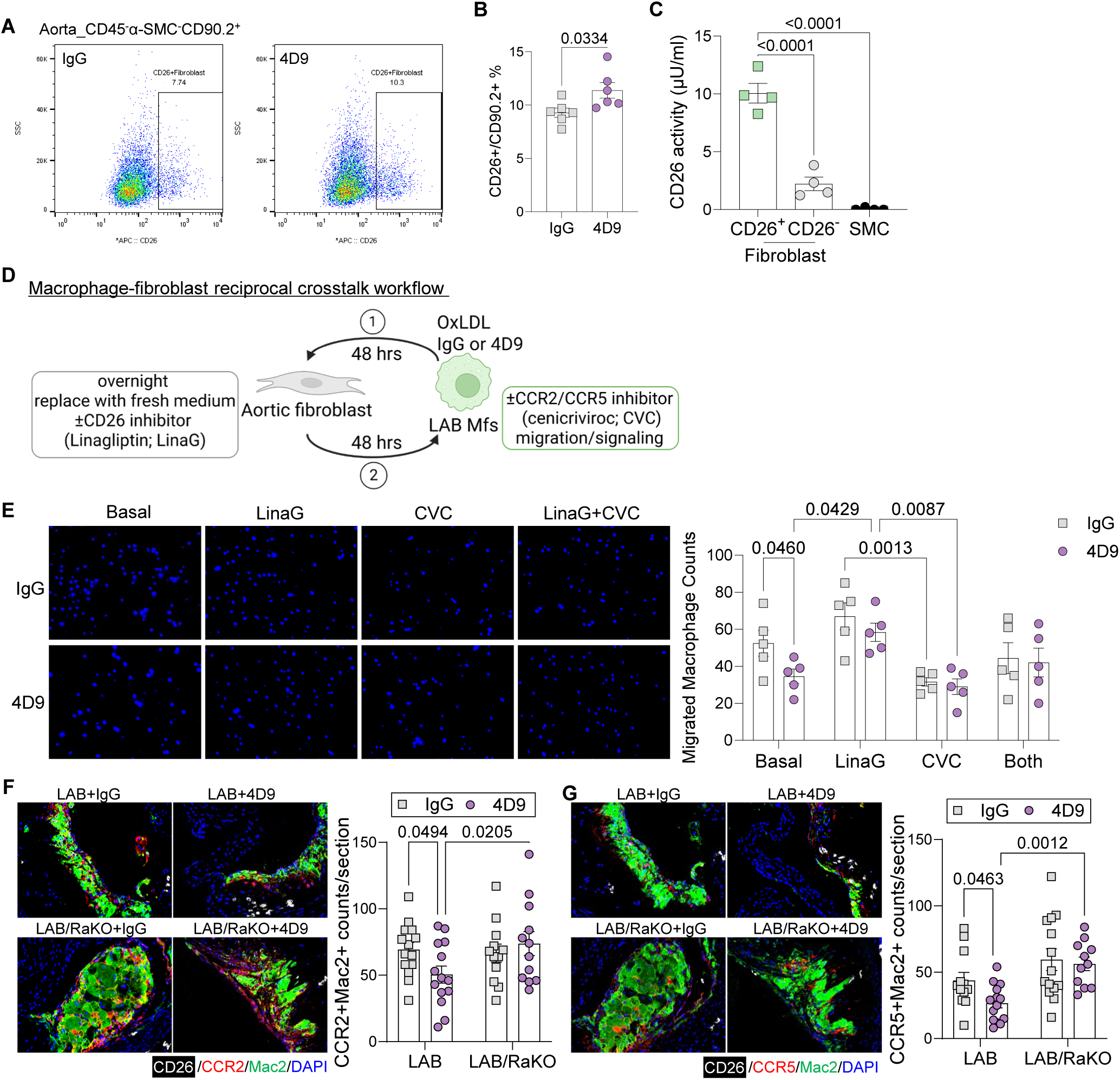
CD26⁺ aortic fibroblasts regulate macrophage recruitment through reciprocal fibroblast-macrophage signaling. **A.** Representative flow cytometry plots showing CD26⁺ fibroblasts within the aorta CD45⁻α-SMC⁻CD90.2⁺ stromal population from atherosclerotic LAB mice treated with IgG or 4D9. **B.** Quantification of CD26⁺ cells as a percentage of CD45⁻α-SMC⁻CD90.2⁺ fibroblasts. **C.** DPP4/CD26 enzymatic activity in sorted aortic CD26⁺ fibroblasts, CD26⁻ fibroblasts, and smooth muscle cells (SMCs). **D.** Schematic of the reciprocal macrophage-fibroblast crosstalk experiments. LAB macrophages were treated with OxLDL (50µg/ml) together with IgG or 4D9 (20μg/ml) for 48 hours, and macrophage-conditioned medium was subsequently transferred to aortic fibroblasts for overnight treatment. The conditioned medium was then removed and replaced with fresh medium with or without the DPP4/CD26 inhibitor linagliptin (LinaG). After an additional 48 h, fibroblast-conditioned medium was collected and used for downstream macrophage migration and signaling assays in the presence or absence of the CCR2/CCR5 inhibitor cenicriviroc (CVC). **E.** Representative images and quantification of macrophage transwell migration in response to fibroblast-conditioned medium under basal conditions or following LinaG, CVC, or combined LinaG+CVC treatment. Nuclei are shown in blue. **F-G.** Representative aortic lesion images and quantification of CCR2⁺Mac2⁺ and CCR5⁺Mac2⁺ macrophages in LAB and LAB/PDGFRaKO (LAB/RaKO) mice treated with IgG or 4D9.

To test whether this fibroblast state could influence macrophage migration, we developed a sequential macrophage-fibroblast conditioned-media system (Fig. 6D). LAB macrophages were treated with OxLDL together with IgG or 4D9 for 48 hours, and macrophage-conditioned medium was transferred to primary aortic fibroblasts. After overnight exposure, the medium was replaced with fresh medium with or without the DPP4 inhibitor linagliptin (LinaG). Fibroblast-conditioned medium was subsequently collected and used for macrophage migration assays in the presence or absence of the CCR2/CCR5 inhibitor Cenicriviroc.

Under basal conditions, fibroblast-conditioned medium generated downstream of 4D9-treated macrophages induced fewer migrating macrophages than medium generated from IgG-treated macrophages (Fig. 6E). Inhibition of fibroblast DPP4 activity with linagliptin increased macrophage migration in the 4D9 condition and eliminated the difference between IgG and 4D9-derived fibroblast-conditioned media (Fig. 6E). In contrast, CCR2/CCR5 blockade with cenicriviroc reduced macrophage migration, supporting a contribution of chemokine-receptor signaling to the migratory response (Fig. 6E). These findings indicate that the reduced macrophage chemotactic activity associated with the 4D9-induced fibroblast state depends, at least in part, on CD26/DPP4 activity.

We next examined whether this relationship was also evident in vivo. Under 4D9 treatment, lesional CCR2^+^Mac2^+^ macrophages were significantly reduced in LAB lesions compared with IgG, whereas this reduction was attenuated in LAB/RaKO lesions, resulting in significantly more CCR2^+^Mac2^+^ macrophages in 4D9-treated LAB/RaKO than LAB lesions (Fig. 6F). A similar pattern was observed for CCR5^+^Mac2^+^ macrophages (Fig. 6G). This correlates with the greatest change in fibroblast to macrophage signaling involving these receptors in the CellChat analysis of scRNA-seq data (Fig 4E). Thus, disruption of the PDGFRα-dependent fibroblast response was associated with increased accumulation of chemokine receptor-positive macrophages within the plaque. Together, these findings support a reciprocal signaling model in which TREM2 activation promotes macrophage PDGFB-PDGFRα-dependent expansion and activation of matrix-associated fibroblasts, including a CD26/DPP4-high population. In addition to contributing to plaque matrix remodeling, this fibroblast state modifies the local chemotactic environment and restrains CCR2/CCR5-dependent macrophage recruitment.

## Discussion

In this study, we identify a bidirectional macrophage-fibroblast signaling pathway downstream of TREM2 activation that coordinates lesional fibroblast responses and macrophage recruitment. Our previous study demonstrated that LAB *Jak2^V617F^* CH impaired macrophage TREM2 expression and that agonistic TREM2 activation with 4D9 increased fibrous cap formation, macrophage PDGFB expression, and the accumulation of PDGFRα+ fibroblast within the lesion and fibrous cap^13^. These findings suggested a macrophage PDGFB-fibroblast PDGFRα pathway but did not establish whether either component was required for the plaque response. Here, complementary conditional deletion of *Pdgfb* in CX3CR1+ monocytes/macrophages and *Pdgfra* in Col1a2-lineage fibroblast-enriched stromal cells establishes the functional requirement for both sides of this signaling axis. We further identify distinct matrix-associated fibroblast populations downstream of this pathway particularly Fmod^+^ and CD26^+^ fibroblast, notably we showed that that CD26^+^ fibroblasts have high DPP4 enzymatic activity, reduce macrophage chemotaxis and may suppress the accumulation of CCR2^+^ and CCR5^+^ macrophages in lesions. Our findings establish a genetically supported macrophage-fibroblast circuit in which TREM2-driven macrophage PDGFB engages stromal PDGFRα to coordinate lesional fibroblast responses, extracellular matrix organization, and fibrous cap formation. These studies show for the first time bidirectional signaling between plaque macrophages and stromal cells that is coordinated by macrophage TREM2 and PDGFB production which signals via PDGFRA in the CD26/DPP4 fibroblast subpopulation to suppress chemokine back-chat and inflammatory macrophage accumulation in plaques. Therapeutic exploitation of this macrophage-stromal cell crosstalk via TREM2 agonism or other approaches involving PDGFB or fibroblast PDGFRA should be particularly effective in the context of inflammatory atherosclerosis where signaling by IL-1β or other factors may act to suppress this beneficial crosstalk at multiple steps.

Several recent studies have highlighted a complex role for TREM2-expressing macrophages in atherosclerosis, with most work focused on lesion development and macrophage burden. Our findings provide a novel explanation for this distinction by showing that TREM2 activation can engage a reparative stromal response through macrophage-derived PDGFB. Thus, the consequences of TREM2 activation are not restricted to macrophage-intrinsic lipid metabolism, survival, or efferocytosis but extend to the regulation of neighboring stromal populations. The interaction is particularly relevant in CH, where inflammatory crosstalk from *Jak2^V617F^* mutant cells suppresses TREM2 and promotes features of plaque instability^13^. Independently, suppression of IL-1β signaling in *Jak2^V617F^* and *Tet2* CH increased fibroblast-like cells within lesions and fibrous cap thickness, whereas depletion of these cells attenuated cap formation^7^. Together, these observations suggest that inflammatory pathways associated with CH may destabilize plaques not only through direct effects on macrophages but also by disrupting beneficial communication between macrophages and matrix-producing stromal populations.

PDGF signaling has long been linked to mesenchymal cell accumulation and fibrous cap formation in atherosclerosis^21^. Lineage-tracing and single-cell studies have further revealed multiple mesenchymal cell states within advanced plaque^25^. In addition to phenotypically modulated SMCs, non-SMC and adventitial-derived populations can contribute fibroblast-like and SMC-like cells, while arterial fibroblasts comprise transcriptionally distinct populations with different matrix-associated programs^25^. Thus, the responses to PDGF in advanced plaques are likely determined not only by ligand availability, but also by the identity and state of the responding cells. Our findings identify a PDGFRα-dependent fibroblast response to macrophage-derived PDGFB. Loss of the 4D9-induced fibrous cap response following either *Pdgfb* deletion in the monocyte/macrophage lineage or *Pdgfra* deletion in Col1a2-lineage stromal cells identifies a macrophage PDGFB-fibroblast PDGFRα pathway distinct from the previously emphasized PDGFRβ-SMC axis. Single-cell analysis further resolved this response across heterogeneous fibroblast populations. Fmod^+^ fibroblasts constituted the major fibroblast population and expanded with 4D9 treatment. Fibromodulin is an extracellular matrix proteoglycan that regulates collagen fibril organization, supporting an association of this population with plaque matrix organization^26^. CD26/Dpp4^+^ fibroblasts represented a smaller population but showed a more pronounced PDGFRα-dependent response, with increased representation and stronger extracellular matrix and collagen-associated programs following TREM2 activation. Thus, TREM2-PDGFB signaling acts across distinct fibroblast states, with CD26^+^ fibroblasts showing a particularly strong dependence on PDGFRα signaling.

CD26^+^ fibroblasts emerged as a particularly distinct stromal population. Recent multimodal single-cell profiling by Bashore et al. identified CD26 as a marker of an arterial fibroblast subset that resides predominantly in the adventitia at baseline but expands into the intima and fibrous cap during atherosclerosis progression^22^. Beyond their matrix-producing functions, fibroblasts are increasingly recognized as regulators of macrophage recruitment through chemokine signaling^27^. Our single-cell analyses showed that CD26^+^ fibroblasts exhibited enhanced extracellular matrix production, collagen organization, and matrix-associated programs, while also displaying prominent predicted communication with lesional macrophages. Cellchat analysis identified *Ccl2/Ccl11-Ccr2/Ccr5* signaling as candidate components of this fibroblast to macrophage interaction. CD26^+^ aortic fibroblasts displayed high DPP4 enzymatic activity, and inhibition of DPP4 with linagliptin increased macrophage migration, whereas CCR2/CCR5 blockade with cenicriviroc reduced migration. Consistent with this interaction in vivo, 4D9 reduced CCR2^+^ and CCR5^+^ macrophages in LAB lesions, an effect attenuated by stromal *Pdgfra* deletion. Thus, our findings suggest CD26^+^ fibroblasts as a PDGFRα-responsive stromal population that links matrix-associated functions with regulation of macrophage recruitment in atherosclerotic plaques.

In summary, our findings establish the causal requirement for a macrophage PDGFB-stromal PDGFRα pathway downstream of TREM2 activation and identify fibroblast heterogeneity as an important component of this response. The results connect macrophage TREM2 signaling to both extracellular matrix remodeling and control of inflammatory-cell recruitment through a CD26/DPP4-dependent fibroblast program. By revealing reciprocal rather than unidirectional communication between macrophages and fibroblasts, these findings provide a cellular mechanism through which TREM2 activation can coordinate inflammatory resolution with fibrous cap remodeling and extend the concept of fibroblasts from passive matrix-producing cells to active regulators of the plaque immune microenvironment. The delineation of this macrophage-matrix cell signaling pathway under the control of macrophage TREM2 agonism provides a framework to understand the potential beneficial effects of TREM2 agonizing antibodies or small molecules as a treatment in advanced inflammatory atherosclerosis as occurs in clonal hematopoiesis.

## Methods

### Sex as a biological variable

Female mice were used for the atherosclerosis studies because they develop advanced atherosclerotic lesions with robust fibrous cap formation in the *Ldlr*^−/−^ model. Similar features of plaque instability have previously been observed in male and female *Jak2^V617F^* clonal hematopoiesis mice.

### Mice

Wild-type C57BL/6J (Strain #: 000664), GFP (C57BL/6-Tg(CAG-EGFP)131Osb/LeySopJ (Strain #: 006567)) *Ldlr^-/-^* (Strain #: 002207), *PDGFRα^fl^* mice (Strain #: 006492), *Col1a2^CreER^* (Strain #:029567) and *Cx3cr1^CreER^* (Strain #:020940) mice were purchased from Jackson Laboratory. *Jak2^V617F^* mice were created and reported previously^28^. *PDGFB^fl^* mice (Stain #: 017622) were kindly provided by Dr. Jiekang Wang and Dr. Mei Wan^29^. *Ldlr^−/−^* recipient mice were fed a western diet (WD) (Harlan Teklad, TD88137) for 16 weeks. In the TREM2 activating antibody (4D9) treated cohort, isotype control IgG (DC1887) and 4D9 (DC1847) mouse antibodies were injected intraperitoneally into the mice at 10 mg/kg per week for 14 weeks. All mice housed under 12 hour’s light/dark cycle and expanded in a pathogen free condition. All protocols were approved by the Institution Animal Care and Use Committee of Columbia.

### Complete blood count

Whole blood was collected into EDTA-coated tubes. White blood cell, red blood cell, and platelet counts were measured using a FORCYTE Veterinary Hematology Analyzer (Oxford Science).

### Atherosclerotic lesion analysis

Hearts were fixed and embedded in paraffin, and serial sections through the aortic root were obtained. Six sections per mouse were stained with H&E for measurement of total lesion and necrotic core areas. Necrotic core was defined as the acellular region within the plaque and quantified using ImageJ. Fibrous cap area was assessed by Masson’s trichrome staining (MilliporeSigma, HT15). The cap region was defined morphologically at the luminal surface of the lesion, and cap area was expressed relative to total lesion area.

### Picrosirius red staining and collagen quantification

Aortic root sections were stained with Picrosirius red according to the manufacturer’s protocol (Polysciences, 24901). Sections were imaged using identical acquisition settings across experimental groups. Collagen-positive area within the plaque was quantified using ImageJ and expressed as a percentage of total lesion area. Representative bright-field images were also obtained where indicated.

### Immunofluorescence staining

Paraffin-embedded aortic root sections were deparaffinized and rehydrated. Antigen retrieval was performed using citrated-based unmasking solution (Vector, H-3300). Sections were blocked and incubated with primary antibodies overnight at 4°C, followed by the appropriate fluorescent secondary antibodies. Slides were mounted with DAPI-containing mounting medium. The antibodies used for staining included the following: anti-Mac-2 (Cedarlane, CL8942AP, 1:10,000); anti-α-SMA (MilliporeSigma, C6198 1:300); anti-PDGFRα (R&D Systems, AF1062, 1:100); anti-CD26 (Abcam, ab187048, 1:100); anti-FMOD (MilliporeSigma, sc-166406, 1:50); anti-CCR5 (R&D systems, NBP2-31374, 1:100); and anti-CCR2 (R&D systems, NBP1-48337, 1:100). Sections were mounted using DAPI (Thermofisher, P36936) and imaged using a Leica DMI6000B microscope. For the FMOD staining, the Vector Laboratories Mouse on Mouse (M.O.M., BMK2202) immunodetection Kit was used to minimize background from the mouse anti-FMOD primary antibody in mouse tissue sections. Image analysis was performed using ImageJ.

### Bone marrow-derived macrophages

Bone marrow cells were isolated from 8-16 weeks mice and differentiated into macrophages in DMEM containing 10% heat-inactivated FBS, penicillin/streptomycin, and 20% L929-conditioned medium for 5-7 days.

### Isolation and culture of primary aortic fibroblasts

Aortas were minced and digested at 37°C in an enzyme cocktail Liberase (Milliporesigma, 5401127001, 1.5mg), hyaluronidase (Milliporesigma, H3506 120 U/mL), and DNase I (Milliporesigma, DN25, 120 U/mL) with gentle agitation. The resulting cell suspension was filtered through a 70-µm cell strainer and washed before downstream flow cytometry or cell sorting. Antibody cocktails including anti-CD45 (BioLegend, 103128, 1:200), anti-CD11b (BioLegend, 101216, 1:200), anti-CD90.2 (Invitrogen,12-0902-82, 1:200), anti-CD26 (BioLegend,137807, 1:100), anti-aSMA (Invitrogen, 53-9760-80, 1:100), and anti-PDGFRA (Invitrogen, 11-1401-82, 1:100). Flow cytometry was performed using BD Fortessa or LSRII or Influx cell sorter, and data were analyzed with FlowJo software (BD).

### Human iPSC-derived macrophages

Isogenic WT and JAK2V617F human iPSCs were differentiated through CD34+ hematopoietic progenitors into macrophages as described previously (). Macrophages were treated with control antibody or agonistic human TREM2 antibody at 20 μg/mL for 48 hours. Cells were collected for RNA and western blot analysis. Cells were lysed in RIPA buffer containing protease and phosphatase inhibitors. Protein concentrations were measured by BCA assay. Equal amounts of protein were mixed with Laemmli sample buffer, heated at 95°C for 5 minutes, separated by SDS-PAGE, and transferred to nitrocellulose membranes. Membranes were blocked in 5% nonfat milk in TBST and incubated with PDGFB primary antibodies (Abcam, ab23914,1:1,000) overnight at 4°C, followed by HRP-conjugated secondary antibodies.

### Quantitative PCR

RNA was isolated using the RNeasy Mini Kit (QIAGEN, 74106) and quantified using a NanoDrop spectrophotometer. cDNA was generated from equal amounts of RNA, and quantitative PCR was performed using SYBR Green Master Mix on an Applied Biosystems real-time PCR system.

### Other reagents and functional assays

Linagliptin (MCE, HY-10284), a PDGFR inhibitor (Cayman, CP673451), and cenicriviroc (Cayman, 23927) were used to inhibit DPP4/CD26, PDGFRα, and CCR2/CCR5 signaling, respectively. Mitomycin C (Cayman, 50-07-7) was used to inhibit fibroblast proliferation during scratching assays. DPP4/CD26 enzymatic activity was measured using a DPP4 Activity Assay Kit (Abcam, ab204722) according to the manufacturer’s instructions. Fibroblast proliferation was assessed by EdU incorporation (Thermofisher, C10639) according to the manufacturer’s instructions. CCL2 Elisa kit (Thermofisher, 88-7391-22) and CCL11 ELISA kit (Protein tech, KE10043). OxLDL (Thermofisher, L34357).

### scRNA-seq analysis

FASTQ files of the multiplexed single-cell RNA sequencing data for the aortic cells were processed with Cell Ranger version 6.1.2. The mm10-2020-A reference transcriptome (GENCODE vM23/Ensembl98) was used to align the sequencing reads, and cells were assigned to their respective samples of origin using four unique hashtag oligonucleotides (HTOs). Seurat 4.3.0 ^30^ in R software version 4.1.0 was then utilized to analyze the unique molecular identifier (UMI) matrix for each sample. Cells expressing fewer than two hundred genes and containing more than ten percent mitochondrial genes, as well as genes expressed in fewer than ten cells, were excluded. For each sample, additional preprocessing steps were carried out according to sequencing depth, with the criteria detailed in Supplemental Table. In each dataset, gene counts were normalized to the total number of genes per cell, then were scaled by a factor of 10,000 and subjected to a natural log transformation. The top 1,000 variable genes were selected from each single-cell sample. The four datasets were then integrated separately for aortic cells, utilizing the top 2,000 highly variable genes identified across the datasets and 20 dimensions of integration anchors. In the examination of aortic single-cell RNA sequencing data, the first 15 principal components (PCs) and the 20 nearest neighbors were used to develop the Shared Nearest Neighbor (SNN) graph. A resolution setting of 0.5 was then utilized through the Louvain clustering to categorize the various cell types. Differential gene expression analysis was performed through the Wilcoxon Rank Sum test to identify the most significantly expressed genes by contrasting one cluster against all others. Genes that were expressed in over 25% of cells, exhibited a minimum fold change of 1.5, and achieved a Bonferroni corrected p-value below 0.05 were classified as differentially expressed for each cell type. Ligand-receptor interaction analysis was performed using the CellChat R package. Ligand-receptor interactions were inferred based on ligand and receptor expression across macrophages, SMCs, and fibroblast populations using the CellChatDB mouse database. Fibroblasts were analyzed as sender populations and myeloid cells as receiver populations to identify potential fibroblast-derived signaling interactions. Communication probabilities were calculated using the CellChat framework, and predicted ligand-receptor interactions were visualized using CellChat.

### Statistics

Data are presented as mean ± SEM unless otherwise indicated. Comparisons involving genotype and antibody treatment were analyzed using 2-way ANOVA with the multiple-comparison test. Two-group comparisons were performed using 2-tailed Student’s t test when data were normally distributed or the Mann-Whitney U test when appropriate. For experiments involving more than 2 groups, 1-way ANOVA followed by the appropriate multiple-comparison test was used. A p value less than 0.05 was considered statistically significant. Statistical analyses were performed using GraphPad Prism. Single-cell enrichment analyses used adjusted P values as indicated.

### Study approval

All animal procedures were approved by the Institutional Animal Care and Use Committee of Columbia University and were performed in accordance with institutional guidelines.

## Data availability

Values for individual data points associated with the manuscript and supplemental figures will be provided in the Supporting Data Values file. Single-cell RNA-sequencing data will be deposited in the NCBI Gene Expression Omnibus, and the accession number will be provided before publication.

## Author contributions

W.L. designed and performed experiments, analyzed data and wrote the manuscript. C.X. designed and conducted scRNA-seq analysis. M.Y., C.H., and T.X. performed the experiments and edited the manuscript. K.S. and C.H. generated and provided IgG and 4D9 antibodies, designed the experiments and edited the manuscript. M.P.R., I.T., N.W. shared the reagents, designed experiments and provided scientific feedback about the manuscript. A.R.T. designed experiments, supervised and directed the project.

---

**Supplemental Fig. 1.**
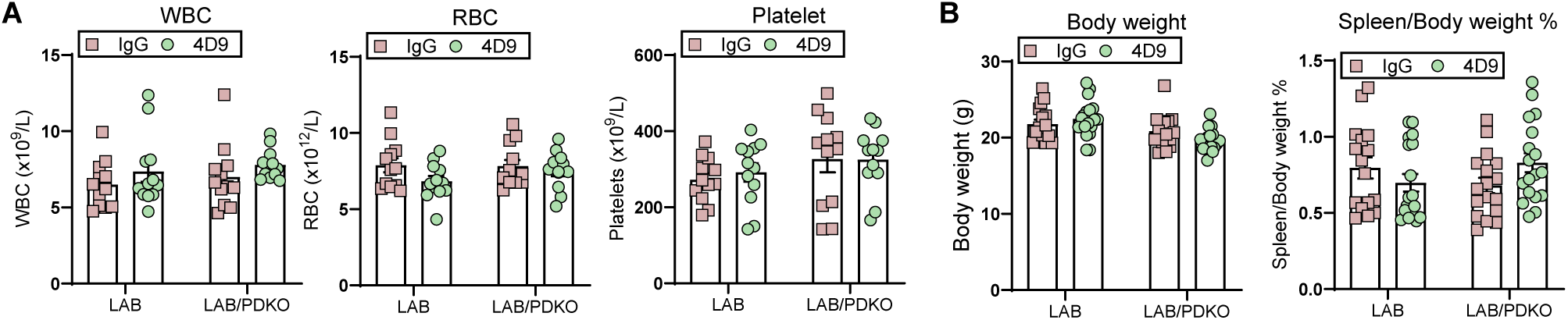
Baseline characterization of LAB/PDGFB KO mice. **A.** Complete blood count analysis showing white blood cell (WBC), red blood cell (RBC) and platelet count in LAB and LAB/PDGFB KO (LAB/PDKO) mice treated with IgG or 4D9. **B.** Body weight and spleen/body weight ratio in the same cohort.

**Supplemental Fig. 2.**
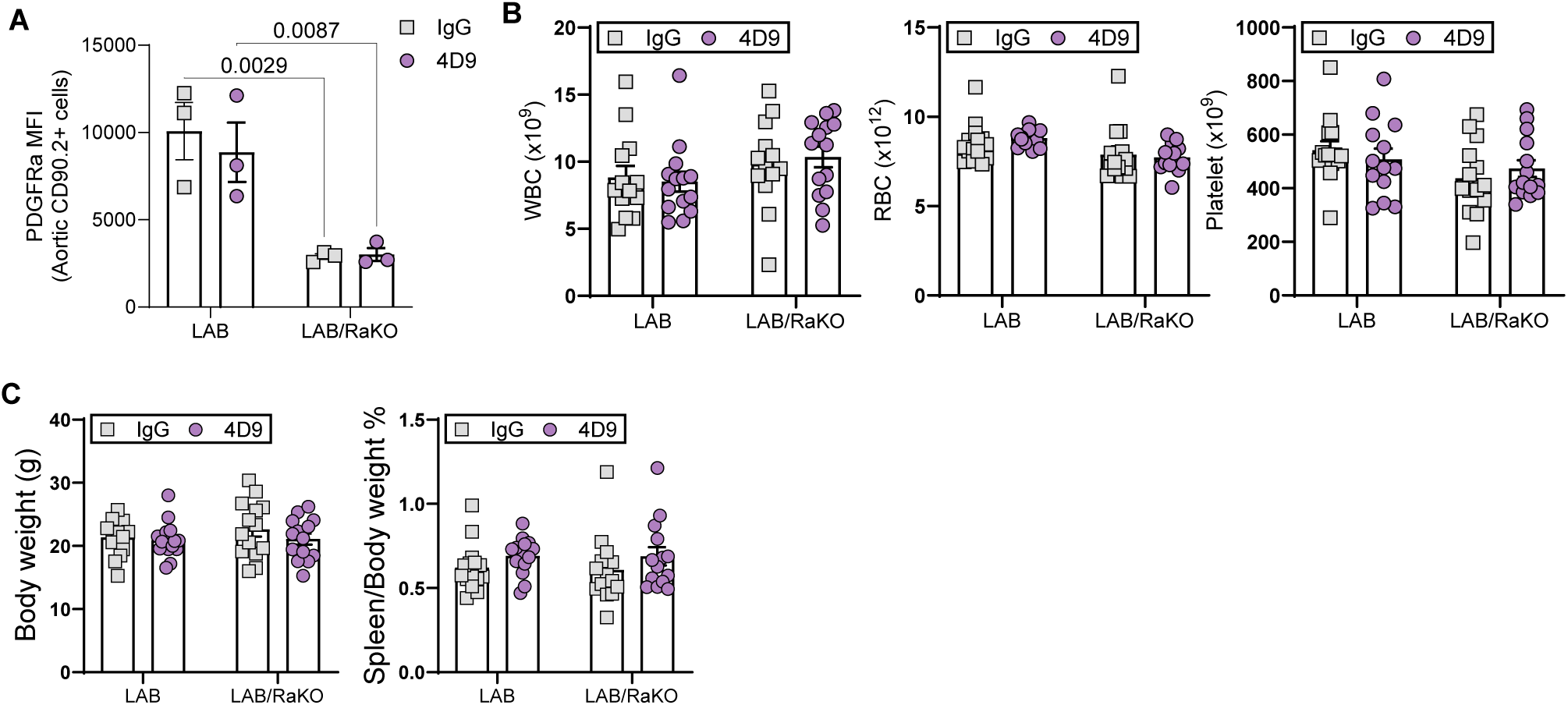
PDGFRa knockout verification and baseline characterization of LAB/PDGFRa mice. **A. B.** Complete blood count analysis showing white blood cell (WBC), red blood cell (RBC) and platelet count in LAB and LAB/PDGFRa (LAB/Ra) mice treated with IgG or 4D9. **C.** Body weight and spleen/body weight ratio in the same cohort.

**Supplemental Fig 3.**
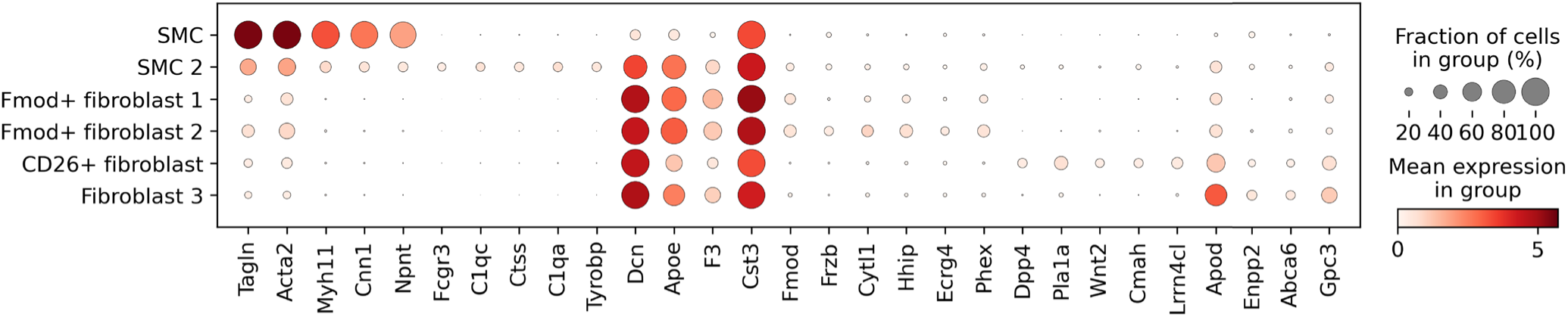
Transcriptional annotation of aortic stromal cell subclusters. Dot plot of selected canonical and subcluster-enriched marker genes used to annotate stromal cell populations. Dot size indicates the fraction of cells within each cluster expressing the indicated gene, and color intensity indicates mean expression.

**Supplemental Fig 4.**
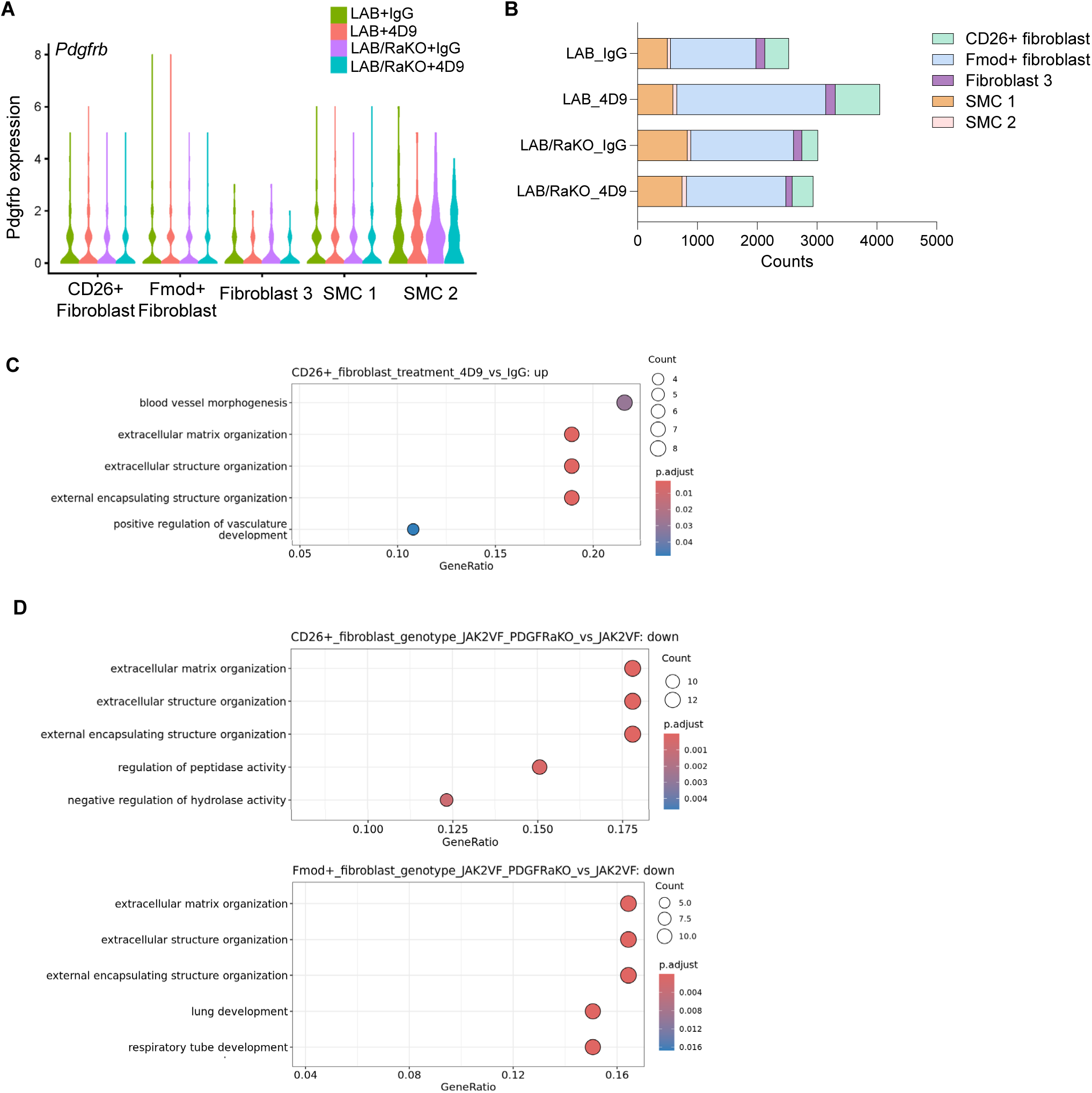
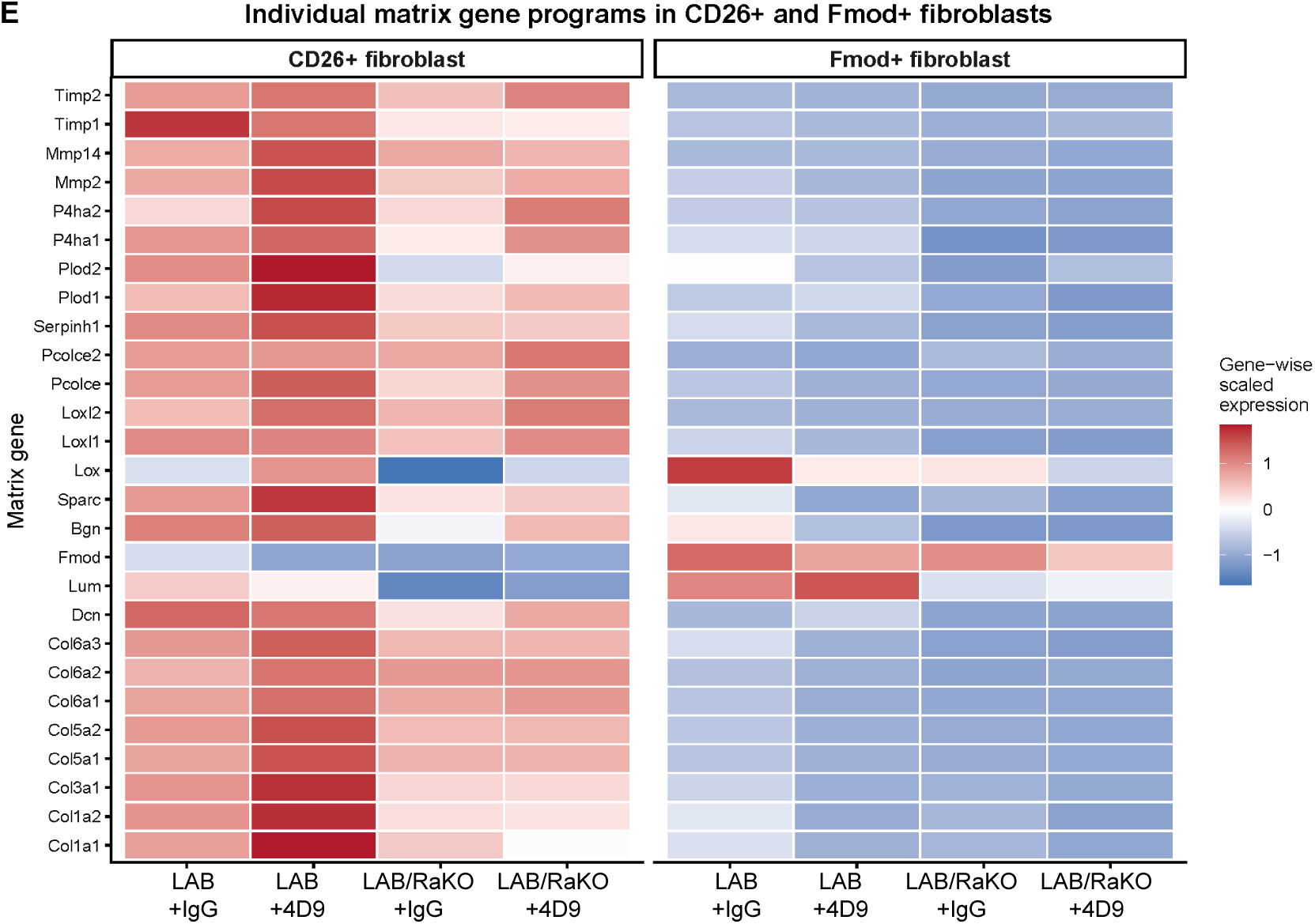
PDGFRα signaling regulates extracellular matrix programs in aortic fibroblast populations. **A.** Violin plots showing log-normalized *Pdgfrb* expression across CD26^+^ fibroblasts, Fmod^+^ fibroblasts, fibroblast 2, SMC1, and SMC2 populations from LAB and LAB/RaKO mice treated with IgG or 4D9. **B.** Number of recovered cells in each annotated stromal population from LAB and LAB/RaKO aortas treated with IgG or 4D9, based on single-cell RNA-sequencing data. **C.** Gene Ontology (GO) enriched among genes upregulated in CD26+ fibroblasts from 4D9-treated compared with IgG-treated LAB mice, including extracellular matrix and vascular organization programs. **D.** GO terms enriched among genes downregulated in Fmod^+^ fibroblasts following stromal *Pdgfra* deletion. Lower, GO terms enriched among genes downregulated in CD26^+^ fibroblasts following stromal *Pdgfra* deletion, including extracellular matrix organization and extracellular structure organization. Dot size represents the number of genes contributing to each GO term, and color indicates the adjusted P value. **E.** Heatmap showing gene-wise scaled expression of individual extracellular matrix and collagen-associated genes in CD26^+^ and Fmod^+^ fibroblasts across LAB+IgG, LAB+4D9, LAB/RaKO+IgG, and LAB/RaKO+4D9 groups. Expression was averaged within each fibroblast population and experimental group and scaled across groups for each gene. Red indicates relatively higher and blue relatively lower expression.

**Supplemental Figure 5.**
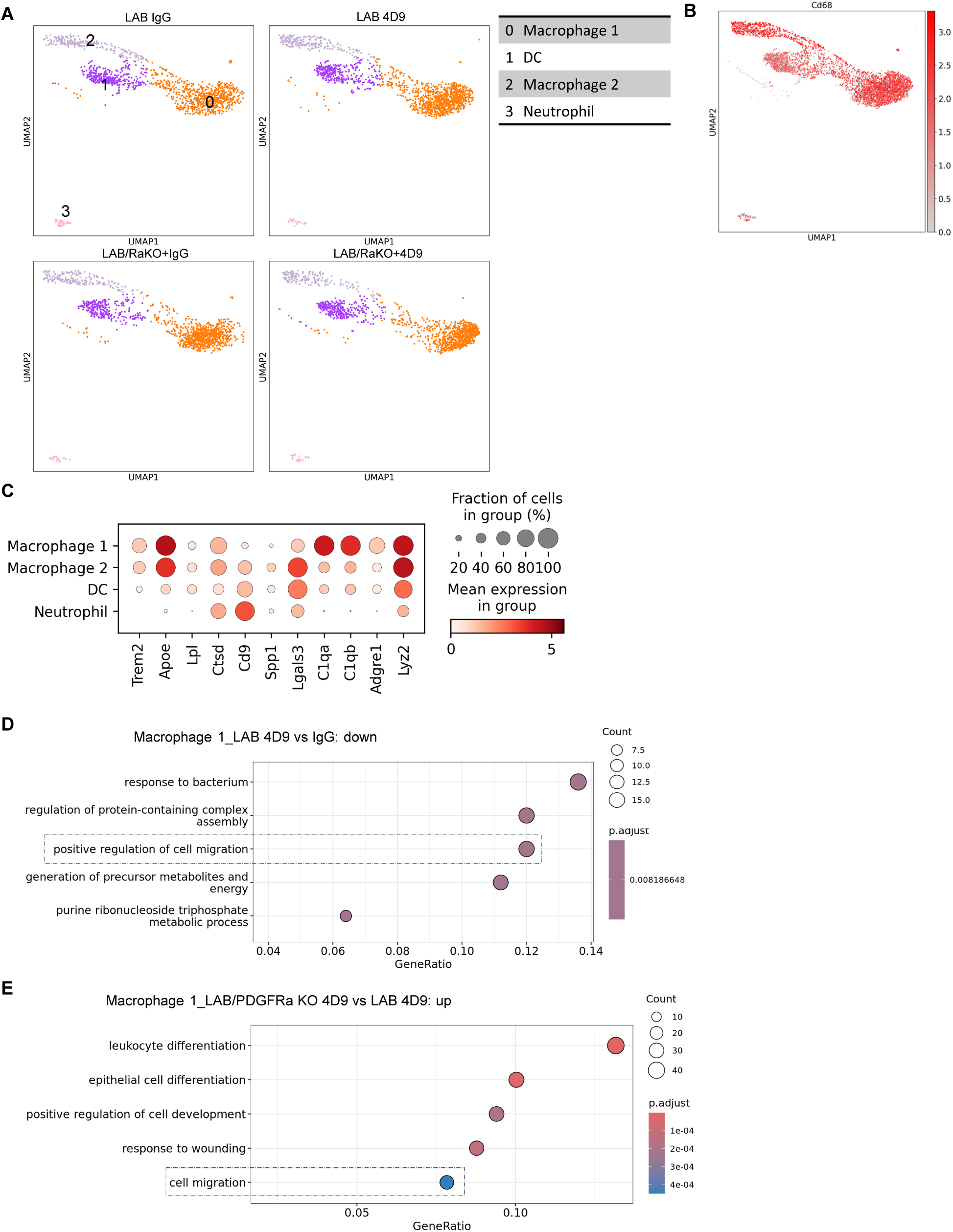
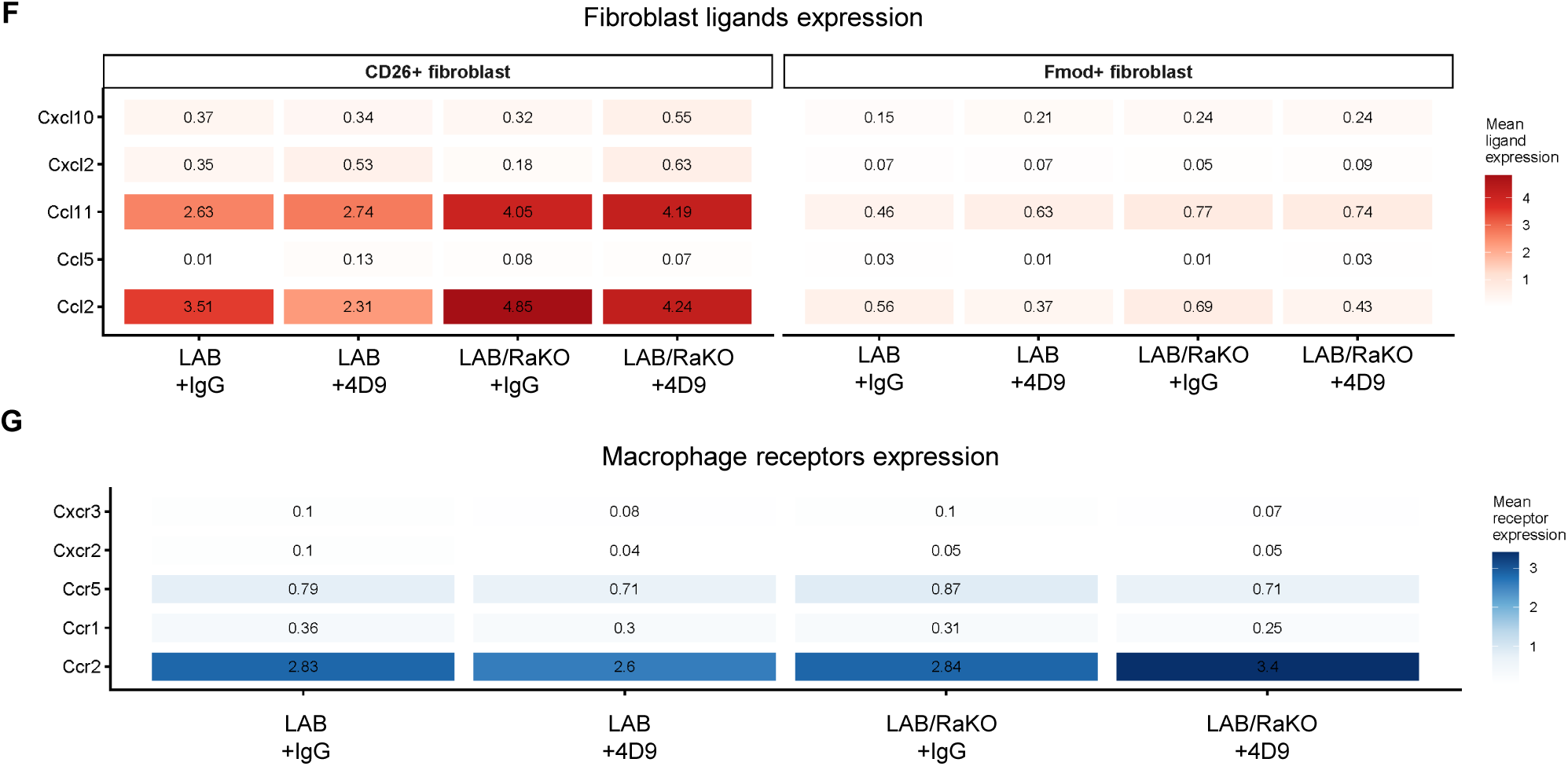
Single-cell analysis identifies macrophage migration programs and fibroblast to macrophage chemokine signaling associated with 4D9 treatment and stromal PDGFRα deficiency. **A.** UMAP visualization of aortic single-cell RNA-seq data from LAB *Jak2^VF^* (LAB) mice treated with IgG or 4D9 and LAB *Jak2^VF^*/PDGFRαKO (LAB/Ra) mice treated with IgG or 4D9. Myeloid cell clusters were identified. Cluster identities are indicated in the accompanying table. **B.** Feature plots showing expression of the macrophage-associated markers *Cd68* and across the integrated aortic cell populations. **C.** Dot plot showing expression of TREM2/macrophage-associated genes, including *Trem2, Apoe, Lpl, Ctsd, Cd9, Spp1, Lgals3, C1qa, C1qb, Adgre1,* and *Lyz2*, across cell clusters. Dot size represents the percentage of cells expressing each gene and color intensity represents average expression. **D.** Gene Ontology (GO) biological process enrichment analysis of genes downregulated in Macrophage 1 from LAB mice treated with 4D9 compared with IgG. Migration-related pathways, including positive regulation of cell migration, are highlighted. **E.** Gene Ontology (GO) biological process enrichment analysis of genes upregulated in Macrophage 1 from LAB/PDGFRαKO mice treated with 4D9 compared with 4D9-treated LAB mice. The migration-related term cell migration is highlighted. Dot size represents gene count and color indicates adjusted *P* value. **F.** Mean expression of selected chemokine ligands in CD26^+^ and Fmod^+^ fibroblasts across 4 groups. Values indicate mean ligand expression within each fibroblast population and treatment group. **G.** Mean expression of corresponding chemokine receptors across the combined macrophage compartment. Values indicate mean receptor expression within the macrophage population for each experimental group.

**Supplemental Figure 6.**
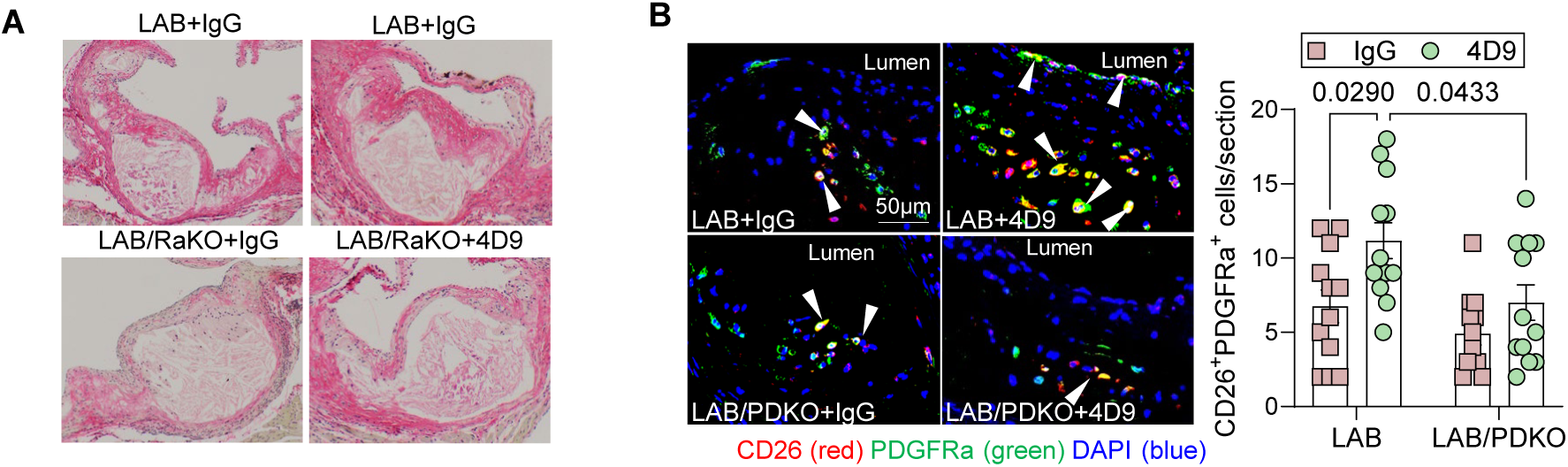
Plaque collagen in the PDGFRα cohorts and histological validation of CD26^+^PDGFRα^+^ fibroblasts in PDGFB cohorts. **A.** Representative bright-field images of Picrosirius red-stained aortic root lesions from LAB and LAB/RaKO mice treated with IgG or 4D9. **B.** Representative immunofluorescence staining of aortic root lesions from LAB and LAB/PDKO mice treated with IgG or 4D9 for CD26 (red), PDGFRα (green), and DAPI (blue). Arrowheads indicate CD26^+^PDGFRα^+^ cells within the plaque and fibrous cap region. Right, quantification of CD26^+^PDGFRα^+^ fibroblast per section.

## References

1. Heron M. Deaths: Leading Causes for 2015. Natl Vital Stat Rep. 2017;66:1–76.

2. Palaniappan LP, Allen NB, Almarzooq ZI, Anderson CAM, Arora P, Avery CL, Baker-Smith CM, Bansal N, Currie ME, Earlie RS, et al. 2026 Heart Disease and Stroke Statistics: A Report of US and Global Data From the American Heart Association. Circulation. 2026;153:e275–e906. doi: 10.1161/CIR.0000000000001412

3. Libby P. The molecular mechanisms of the thrombotic complications of atherosclerosis. J Intern Med. 2008;263:517–527. doi: 10.1111/j.1365-2796.2008.01965.x

4. Sarraju A, Nissen SE. Atherosclerotic plaque stabilization and regression: a review of clinical evidence. Nat Rev Cardiol. 2024;21:487–497. doi: 10.1038/s41569-023-00979-8

5. Wirka RC, Wagh D, Paik DT, Pjanic M, Nguyen T, Miller CL, Kundu R, Nagao M, Coller J, Koyano TK, et al. Atheroprotective roles of smooth muscle cell phenotypic modulation and the TCF21 disease gene as revealed by single-cell analysis. Nat Med. 2019;25:1280–1289. doi: 10.1038/s41591-019-0512-5

6. Bashore AC, Coronel J, Xue C, Zhu LY, Reilly MP. Single-Cell Multimodal Profiling Reveals a Novel CD26(+) Fibroblast Subpopulation in Atherosclerosis-Brief Report. Arterioscler Thromb Vasc Biol. 2025;45:1389–1397. doi: 10.1161/ATVBAHA.124.322370

7. Fidler TP, Dunbar A, Kim E, Hardaway B, Pauli J, Xue C, Abramowicz S, Xiao T, O’Connor K, Sachs N, et al. Suppression of IL-1beta promotes beneficial accumulation of fibroblast-like cells in atherosclerotic plaques in clonal hematopoiesis. Nat Cardiovasc Res. 2024;3:60–75. doi: 10.1038/s44161-023-00405-9

8. Li G, Asselberghs SEJ, Yu B, Feng C, Mendez PL, Kasakovski D, Goossens P, Ackermans TM, Tillie R, Gijbels MJ, et al. Atherosclerotic plaque fibroblasts derive from adventitial and medial Pdgfra-lineage-positive cells and predominantly maintain fibroblast identity. Cardiovasc Res. 2026. doi: 10.1093/cvr/cvag183

9. van Kuijk K, McCracken IR, Tillie R, Asselberghs SEJ, Kheder DA, Muitjens S, Jin H, Taylor RS, Wichers Schreur R, Kuppe C, et al. Human and murine fibroblast single-cell transcriptomics reveals fibroblast clusters are differentially affected by ageing and serum cholesterol. Cardiovasc Res. 2023;119:1509–1523. doi: 10.1093/cvr/cvad016

10. Patterson MT, Firulyova MM, Xu Y, Hillman H, Bishop C, Zhu A, Hickok GH, Schrank PR, Ronayne CE, Caillot Z, et al. Trem2 promotes foamy macrophage lipid uptake and survival in atherosclerosis. Nat Cardiovasc Res. 2023;2:1015–1031. doi: 10.1038/s44161-023-00354-3

11. Patterson MT, Xu Y, Hillman H, Osinski V, Schrank PR, Kennedy AE, Barrow F, Zhu A, Tollison S, Shekhar S, et al. Trem2 Agonist Reprograms Foamy Macrophages to Promote Atherosclerotic Plaque Stability-Brief Report. Arterioscler Thromb Vasc Biol. 2024;44:1646–1657. doi: 10.1161/ATVBAHA.124.320797

12. van Lengerich B, Zhan L, Xia D, Chan D, Joy D, Park JI, Tatarakis D, Calvert M, Hummel S, Lianoglou S, et al. A TREM2-activating antibody with a blood–brain barrier transport vehicle enhances microglial metabolism in Alzheimer’s disease models. Nature Neuroscience. 2023;26:416–429. doi: 10.1038/s41593-022-01240-0

13. Liu W, Hardaway BD, Kim E, Pauli J, Wettich JL, Yalcinkaya M, Hsu CC, Xiao T, Reilly MP, Tabas I, et al. Inflammatory crosstalk impairs phagocytic receptors and aggravates atherosclerosis in clonal hematopoiesis in mice. J Clin Invest. 2024;135. doi: 10.1172/JCI182939

14. Rutherford C, Martin W, Carrier M, Anggard EE, Ferns GA. Endogenously elicited antibodies to platelet derived growth factor-BB and platelet cytosolic protein inhibit aortic lesion development in the cholesterol-fed rabbit. Int J Exp Pathol. 1997;78:21–32. doi: 10.1046/j.1365-2613.1997.d01-237.x

15. Tang J, Kozaki K, Farr AG, Martin PJ, Lindahl P, Betsholtz C, Raines EW. The absence of platelet-derived growth factor-B in circulating cells promotes immune and inflammatory responses in atherosclerosis-prone ApoE-/-mice. Am J Pathol. 2005;167:901–912. doi: 10.1016/S0002-9440(10)62061-5

16. Kozaki K, Kaminski WE, Tang J, Hollenbach S, Lindahl P, Sullivan C, Yu JC, Abe K, Martin PJ, Ross R, et al. Blockade of platelet-derived growth factor or its receptors transiently delays but does not prevent fibrous cap formation in ApoE null mice. Am J Pathol. 2002;161:1395–1407. doi: 10.1016/S0002-9440(10)64415-X

17. Tillie R, Theelen TL, van Kuijk K, Temmerman L, de Bruijn J, Gijbels M, Betsholtz C, Biessen EAL, Sluimer JC. A Switch from Cell-Associated to Soluble PDGF-B Protects against Atherosclerosis, despite Driving Extramedullary Hematopoiesis. Cells. 2021;10. doi: 10.3390/cells10071746

18. Sano H, Sudo T, Yokode M, Murayama T, Kataoka H, Takakura N, Nishikawa S, Nishikawa SI, Kita T. Functional blockade of platelet-derived growth factor receptor-beta but not of receptor-alpha prevents vascular smooth muscle cell accumulation in fibrous cap lesions in apolipoprotein E-deficient mice. Circulation. 2001;103:2955–2960. doi: 10.1161/01.cir.103.24.2955

19. He C, Medley SC, Hu T, Hinsdale ME, Lupu F, Virmani R, Olson LE. PDGFRβ signalling regulates local inflammation and synergizes with hypercholesterolaemia to promote atherosclerosis. Nature Communications. 2015;6:7770. doi: 10.1038/ncomms8770

20. Kim H-J, Cheng P, Travisano S, Weldy C, Monteiro JP, Kundu R, Nguyen T, Sharma D, Shi H, Lin Y, et al. Molecular mechanisms of coronary artery disease risk at the PDGFD locus. Nature Communications. 2023;14:847. doi: 10.1038/s41467-023-36518-9

21. Raines EW. PDGF and cardiovascular disease. Cytokine Growth Factor Rev. 2004;15:237–254. doi: 10.1016/j.cytogfr.2004.03.004

22. Bashore AC, Coronel J, Xue C, Zhu LY, Reilly MP. Single-Cell Multimodal Profiling Reveals a Novel CD26^+^ Fibroblast Subpopulation in AtherosclerosisC#x2014;Brief Report. Arteriosclerosis, Thrombosis, and Vascular Biology. 2025;45:1389–1397. doi: doi:10.1161/ATVBAHA.124.322370

23. Hedlund H, Mengarelli-Widholm S, Heinegard D, Reinholt FP, Svensson O. Fibromodulin distribution and association with collagen. Matrix Biol. 1994;14:227–232. doi: 10.1016/0945-053x(94)90186-4

24. Shami A, Tengryd C, Asciutto G, Bengtsson E, Nilsson J, Hultgardh-Nilsson A, Goncalves I. Expression of fibromodulin in carotid atherosclerotic plaques is associated with diabetes and cerebrovascular events. Atherosclerosis. 2015;241:701–708. doi: 10.1016/j.atherosclerosis.2015.06.023

25. Pan H, Xue C, Auerbach BJ, Fan J, Bashore AC, Cui J, Yang DY, Trignano SB, Liu W, Shi J, et al. Single-Cell Genomics Reveals a Novel Cell State During Smooth Muscle Cell Phenotypic Switching and Potential Therapeutic Targets for Atherosclerosis in Mouse and Human. Circulation. 2020;142:2060–2075. doi: 10.1161/CIRCULATIONAHA.120.048378

26. Shami A, Gustafsson R, Kalamajski S, Krams R, Segers D, Rauch U, Roos G, Nilsson J, Oldberg A, Hultgardh-Nilsson A. Fibromodulin deficiency reduces low-density lipoprotein accumulation in atherosclerotic plaques in apolipoprotein E-null mice. Arterioscler Thromb Vasc Biol. 2013;33:354–361. doi: 10.1161/ATVBAHA.112.300723

27. Tsukui T, Wolters PJ, Sheppard D. Alveolar fibroblast lineage orchestrates lung inflammation and fibrosis. Nature. 2024;631:627–634. doi: 10.1038/s41586-024-07660-1

28. Mullally A, Lane SW, Ball B, Megerdichian C, Okabe R, Al-Shahrour F, Paktinat M, Haydu JE, Housman E, Lord AM, et al. Physiological Jak2V617F expression causes a lethal myeloproliferative neoplasm with differential effects on hematopoietic stem and progenitor cells. Cancer Cell. 2010;17:584–596. doi: 10.1016/j.ccr.2010.05.015

29. Wang J, Fang CL, Noller K, Wei Z, Liu G, Shen K, Song K, Cao X, Wan M. Bone-derived PDGF-BB drives brain vascular calcification in male mice. J Clin Invest. 2023;133. doi: 10.1172/JCI168447

30. Stuart T, Butler A, Hoffman P, Hafemeister C, Papalexi E, Mauck WM, 3rd, Hao Y, Stoeckius M, Smibert P, Satija R. Comprehensive Integration of Single-Cell Data. Cell. 2019;177:1888–1902 e1821. doi: 10.1016/j.cell.2019.05.031

